# High-resolution three-dimensional mapping of eelgrass (*Zostera marina*) habitat and blue carbon using drone-borne LiDAR

**DOI:** 10.1101/2025.08.12.669815

**Authors:** Charles P. Lavin, Toms Buls, Hege Gundersen, Kristina Øie Kvile, Øyvind Tangen Ødegaard, Robert Nøddebo Poulsen, Kasper Hancke

## Abstract

The accessibility of flying drones (Unoccupied Aerial Vehicles) presents scientists and managers with reproducible and cost-effective methods to monitor submerged aquatic vegetation. In particular, drone-borne topobathymetric LiDAR provides high-resolution (cm-scale), three-dimensional information about the geometry and structure of surveyed areas, allowing for quantification of vegetation volume in addition to bathymetry. For habitat-forming submerged and intertidal vegetation like seagrass, this information can advance research regarding the structure and patchiness of canopies in relation to biodiversity, blue carbon storage, and hydrodynamic processes. Here, we report how drone-borne LiDAR can be used to estimate the habitat volume of eelgrass (*Zostera marina*) within a sheltered bay in south-eastern Norway. After classifying LiDAR points using a Random Forest model, we created a Digital Terrain Model of the sea floor and a Digital Surface Model of the eelgrass canopy. From these models, we estimated eelgrass canopy volume to range between 862 and 1099 m^3^ across the small study area. From the volume, we estimated above-ground carbon storage in living eelgrass tissue to range between 96 and 122 kg. To our knowledge, this is the first study to utilise drone-borne LiDAR to quantify the volume and carbon-storage potential of a marine habitat-forming species like eelgrass, thereby demonstrating the potential of drone-borne LiDAR as an efficient tool to provide reproducible and high-resolution data for submerged aquatic habitats, including seagrass meadows.

## Introduction

The ecosystem services provided by seagrasses are numerous, and increasingly important in the context of rapid climatic change and widespread habitat degradation (Duarte and others 2025). Their high primary productivity and ability to trap and retain organic matter show that seagrass sediments are globally important for blue carbon storage (Duarte and others 2005; Fourqurean and others 2012; Röhr and others 2018; Gagnon and others 2024), hence their formal inclusion in 2013 into the IPCC’s Guidelines for Greenhouse Gas Inventories for Wetlands (IPCC 2014). The presence of seagrass beds has also demonstrated enhanced protection from coastal erosion (Infantes and others 2022). As habitat-forming species, seagrasses provide structural complexity that support diverse benthic infauna and epifauna, offering nursery and refuge areas for invertebrates and fishes, as well as sources of food for megaherbivores and predators (McDevitt-Irwin and others 2016; Whitfield 2017; Kindeberg and others 2022; Gagnon and others 2023). These habitat provisions have been shown to support the basis of fisheries productivity globally (Kritzer and others 2016; Unsworth and others 2019; Jänes and others 2021).

Seagrasses are experiencing widespread global decline and degradation (Waycott and others 2009; Dunic and others 2021). In recognition of this, seagrass protection and restoration was included in the Conservation of Migratory Species (CMS) resolution at the 14th United Nations Conference of the Parties (COP14) in 2024 (Unsworth and Jones 2024). This resolution focuses on the need for monitoring of seagrass habitat and associated biodiversity, particularly in relation to known drivers of seagrass loss (Unsworth and Jones 2024). The monitoring of seagrass has traditionally relied on *in-situ* observation and manual data collection, such as snorkel or SCUBA-based surveys. In general, these approaches are often labour-intensive, costly, prone to human error, and restricted to easily accessible areas (Rowan and Kalacska 2021). Conversely, flying drones (technically referred to as Unoccupied Aerial Vehicles, UAVs) have proven effective to expedite a non-invasive, reproducible and efficient alternative for habitat monitoring across a variety of habitats and conditions (Duffy et al., 2018; Joyce et al., 2023). For example, drones mounted with optical sensors have successfully been used to map submerged vegetated habitats including seagrass and macroalgae, by differentiating objects and species based on their unique optical signatures (Gundersen and others 2024; Kvile and others 2024).

Air-borne Light Detection and Ranging (LiDAR) systems carried on small, crewed aircrafts have been applied to map terrestrial and marine ecosystems in three-dimensional space (Chen and others 2023; Borja and others 2024; Lu and Jiang 2024), particularly used in forest research (Mazlan and others 2022). However, drone-mounted systems now offer similar capabilities at lower cost and higher spatial precision, although with a somewhat more limited spatial range (Borja and others 2024). Typically, drone-borne LiDAR collected point cloud densities are at a scale of some hundred points per square meter, while crewed aircrafts collect a point density one magnitude lower.

LiDAR systems work by emitting pulsed laser beams, whereby the time it takes for the laser to reflect off a surface and return to the sensor is used to calculate the distance, and thus the elevation of the targeted surface (Baltsavias 1999). The resulting point cloud data therefore captures the geometry and height of surfaces within the scanned area (Lim and others 2003). Topobathymetric LiDAR surveys land, water, and submerged surfaces, by using a green laser of wavelength 532 nanometres (nm) that penetrates the water column down to 1 – 3 Secchi depths, depending on water clarity and the power of the LiDAR system (Pricope and Bashit 2023). The underwater elevation of points in topobathymetric LiDAR therefore correspond to the elevation of the reflected surface relative to the water surface. This is the case as these systems are optimized to detect changes in the green laser beam signal at the air-water interface. There, part of the laser beam is scattered back to the LiDAR receiver, while other parts of the laser continue to propagate through the water column at slower speeds (Figure 1) (Mandelburger, 2020). In summary, topobathymetric LiDAR generates point clouds that provide detailed elevation profiles of both the sea floor and sea floor cover, i.e. submerged vegetation, facilitating geometric investigation of their structure.

**Figure 1:**
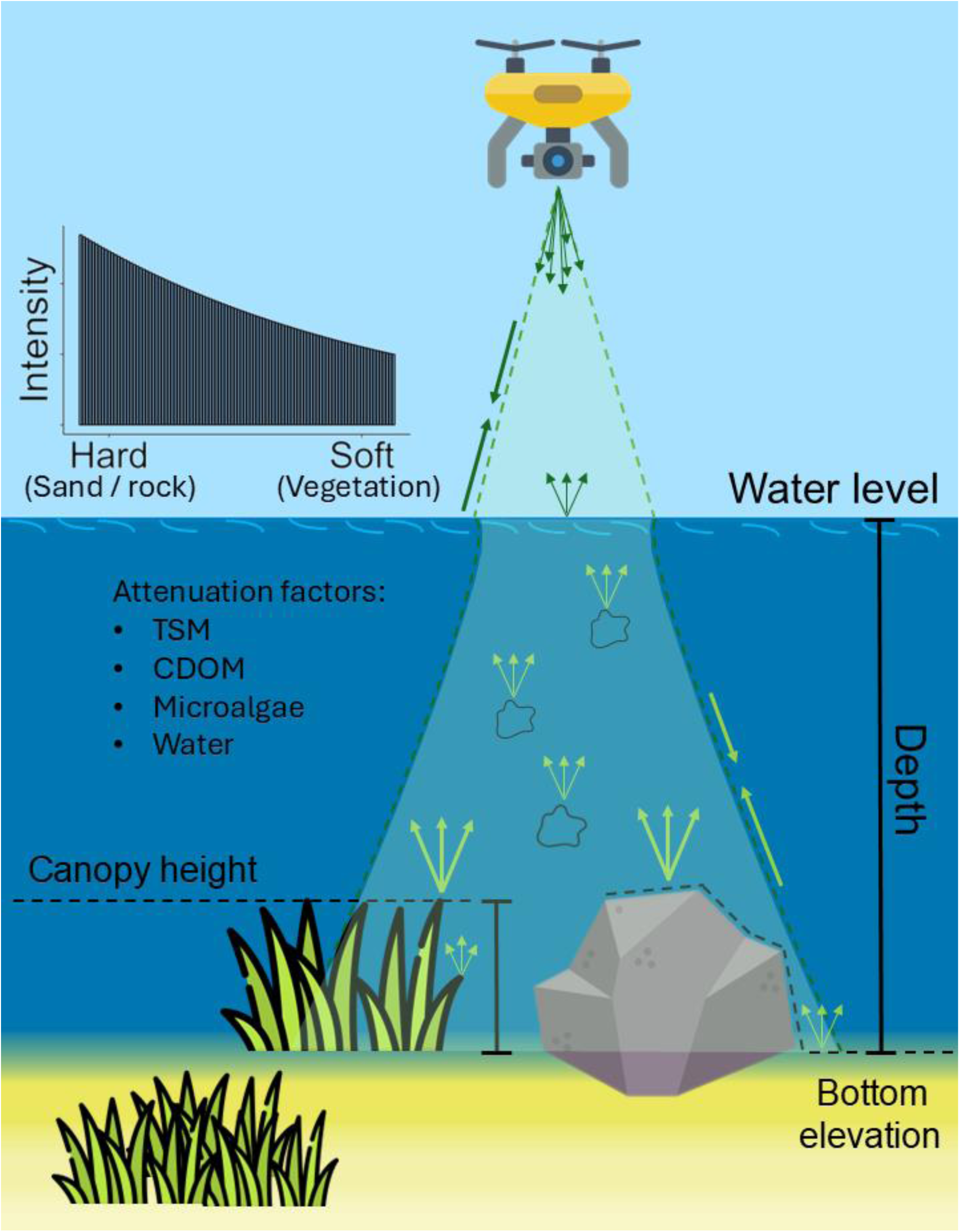
Schematic of drone-borne green topobathymetric LiDAR system. A green laser of wavelength 532 nanometres (nm) is emitted from a drone, whereby at the water surface some of the laser beams are reflected back, whilst the remainder is refracted and continues through the water column. Elevation values are generated based on the time it takes for the laser pulse to return to the sensor and represent elevation values relative to the water surface. Reflections can occur from the sea floor as well as from suspended particles, with maximum depth penetration between 1-3 Secchi depths. Laser attenuation is therefore influenced by water characteristics such as total suspended matter (TSM), coloured dissolved organic matter (CDOM), and the presence of microalgae. By reaching the sea floor, the strength of the reflected signal (i.e. intensity) varies depending on substrate type, allowing distinction between hard (e.g. sand, rock) and soft, absorptive surfaces (i.e. vegetation).

Not only does LiDAR capture geometric information, it also captures radiometric data (Yan and Shaker 2014), including the intensity, or return strength, of the reflected laser beam (Hall and others 2005). The scale of intensity represents the proportion of light reflected from a surface, which varies depending on the material and texture of the target. This variation has been used to differentiate between ground structures and vegetation (Guo and others 2011), and also to distinguish between different species of terrestrial plants (García and others 2010). Overall, the inclusion of intensity values alongside geometric data greatly enhances LiDAR application, especially to distinguish vegetation from natural substrates or artificial surfaces (Wang and Philpot 2007; Yan and others 2012).

Recent advances in drone-borne LiDAR technology have made medium-sized drones increasingly accessible to a wide variety of users, including researchers and managers. However, drone-borne topobathymetric LiDAR systems (i.e., hardware plus software) remain few. Currently, some available systems offer dedicated software for automated post-processing, including georeferencing, water surface detection, and point cloud intensity correction at ‘survey-grade’ quality. Such accessibility of drone technology, paired with user-friendly data handling, can enable cost-effective and reproducible methods necessary to meet global ambitions of seagrass monitoring. Moreover, the geometry of LiDAR data enables three-dimensional quantification and visualisation of surveyed areas at high-resolution, often at scales of a few centimetres. Such data could enhance our understanding of how the structure of seagrass meadows relates to ecological functions, including habitat provisioning, hydrodynamic, and physical coastal conditions, as well as biomass and blue carbon storage capabilities. To the authors’ knowledge, no published studies have yet demonstrated the three-dimensional quantification of seagrass habitat using drone-borne LiDAR.

In the present study, we do so by analysing geometric and radiometric LiDAR data collected by a medium-sized drone (4.1 kg including sensor). Using a Random Forest (RF) model, we classified points within the LiDAR point cloud and created a Digital Terrain Model (DTM) and a Digital Surface Model (DSM) of the sea floor bathymetry and vegetation canopy cover, respectively. Together, these models provide high-resolution volumetric information and three-dimensional visualisation of eelgrass habitat area at a study site in south-eastern Norway, facilitating calculation of aboveground biomass and associated carbon storage.

## Methods

### Study species and area

Eelgrass (*Zostera marina*) is a common seagrass species found throughout the northern hemisphere (Yu and others 2023). In Europe, its distribution spans from sub- tropical conditions in southern Iberia to Arctic conditions in northern Norway (Short and Green 2003). Our study area was located at Ølbergholmen, a small peninsula in Vestfold County, south-eastern Norway (59° N, 10.13° E, Figure 2). The northern section of Ølbergholmen is a sheltered, shallow bay containing eelgrass meadows and thus was chosen as the sampling site.

**Figure 2:**
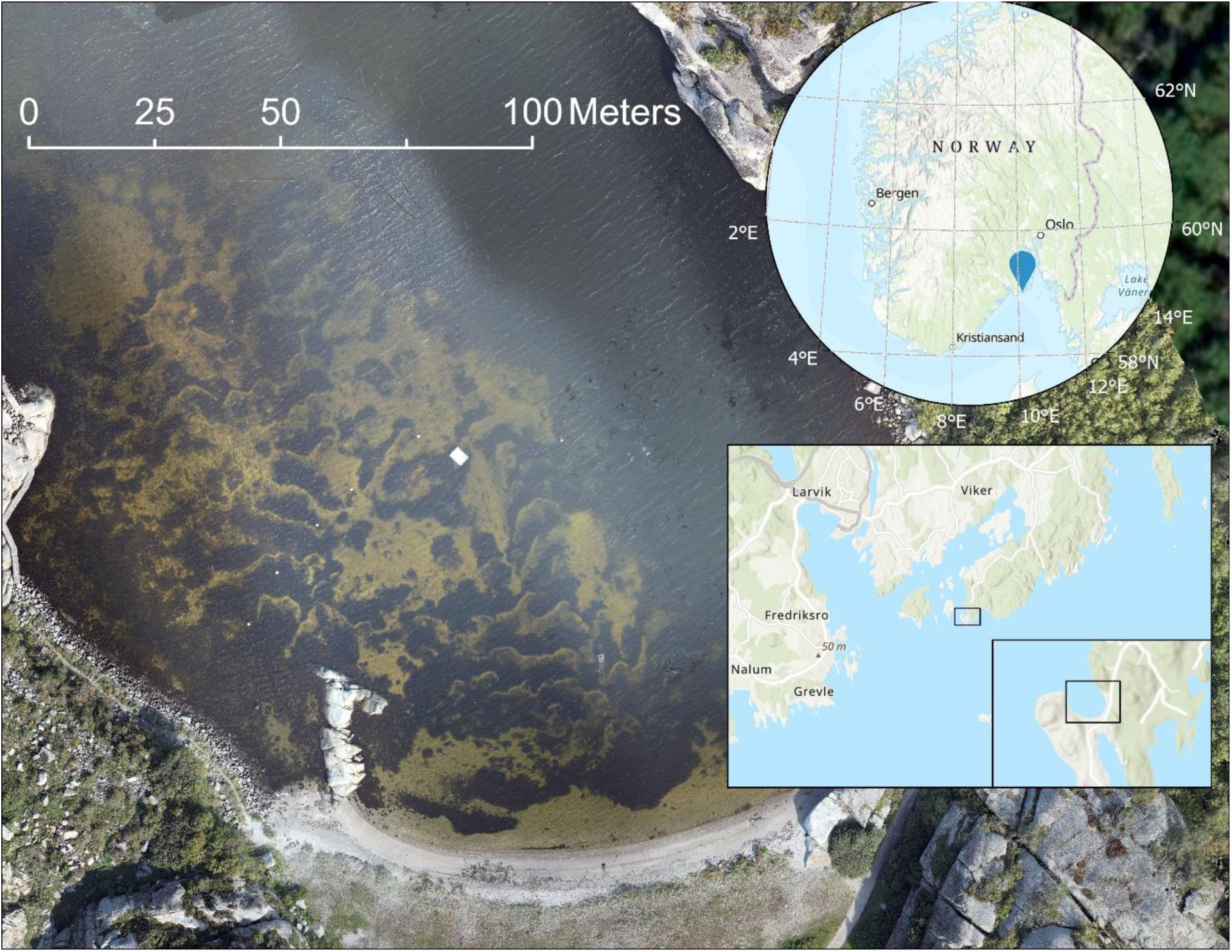
Location of the study site at Ølbergholmen in Vestfold County, Norway, showing a patchy seagrass (*Zostera marina*) meadow on a sandy substrate with scattered macroalgae. Inset maps were created using ArcGIS Pro (ESRI 2025), while the main image is an RGB drone photo taken from 60 m altitude (available from the SeaBee Research Infrastructure’s Geo-Visualization Portal: https://geonode.seabee.sigma2.no).

### LiDAR data collection

Data were collected using a topobathymetric LiDAR sensor (Navigator, YellowScan, France) mounted on a midrange UAV (Tundra 2 equipped with Endurance rotor arms, Hexadrone, France) (YellowScan 2024b). This system includes a green laser with an emitting wavelength of 532 nm and achieves horizontal and vertical accuracies of 0.5 and 2.1 cm, respectively (following calibration in line with the manufacturer’s guidelines). A constant beam angle of 20° to either side of nadir resulted in a 40° field of view for scanning (YellowScan 2024b). The first of two drone flights was conducted on 29 August 2024 at 05:35 hours, at an elevation of 50 m, with 50% overlap between flight lines as recommended by the manufacturer. A second flight occurred later the same day at 06:18 hours, at 25 m elevation, also with 50% overlap. In subsequent analyses, we compare LiDAR-derived eelgrass canopy metrics at both altitudes to assess the effect of drone elevation, and the resulting point cloud density, on canopy volume estimates.

### *In-situ* data collection

To validate LiDAR-generated canopy height and ground elevation values, eelgrass canopy height, and ground elevation both above and below water were collected *in-situ* on 2 November 2024 (Supplementary Figure 1). As *in-situ* measurements were taken after the LiDAR flight, to assess potential seasonal changes in canopy height, we compared canopy measurements from a previous campaign to the same area in September 2023 to data from November 2024. We expected a reduced height in November, which is beyond the period of maximum eelgrass height, which typically occurs around September in the region (Clausen and others 2014). Canopy height was measured visually using mask and snorkel, with height corresponding to the average canopy height (in m) within a circle of 20 cm radius. Ground elevation data were gathered using an Emlid Reach RS3 real-time kinematic (RTK) positioning Global Navigation Satellite System (GNSS) receiver (Emlid 2024) with horizontal and vertical accuracies of 1.2 and 1.3 cm, respectively. Elevation values under water were taken on sandy substrates only (Supplementary Figure 1).

### LiDAR bathymetry comparison with sonar

For deeper areas (> 1.5 m) that could not be accessed with the pole-mounted handheld GNSS, bathymetry data were collected using a BioSonics MX 200 kHz single-beam aquatic habitat mapping echosounder, installed on a Uncrewed Surface Vehicle (USV, Maritime Robotics Otter Pro). This small USV (2 m long, 70 kg) was deployed across the study area using vehicle guidance to follow a pre-programmed mission ‘lawnmower’ pattern, at approximately 1 m/s with 5-10 m cross-track spacing (Supplementary Figure 1), utilising a SGB Ellipse-D Internal Navigation System (INS) and dual GNSS antennas. The echosounder was mounted on the USV’s bow, with a 9° conical beam angle transducer pointed at the sea floor, with the transducer connected to BioSonics MX acquisition electronics contained within the vehicle’s payload. The system logged the transducer signals along with time, latitude, and longitude from the vehicle’s INS. Echosounder data were collected on 13 September 2023 at 08:48 hours, and data processing was performed in BioSonics’ Visual Aquatic software (BioSonics 2025).

### Raw LiDAR data post-processing

LiDAR data was refined following the flight trajectory and georeferenced using YellowScan’s CloudStation software (YellowScan 2024a). This was done by accessing GNSS base stations to account for drone position and altitude, plus an Inertial Measurement Unit (IMU) contained in the LiDAR sensor that accounts for scanner range and angle (Shan and Toth 2018). The resulting point cloud was georeferenced to the local coordinate system (UTM Zone 32N, ETRS 1989) and vertical reference frame (NN2000) (Lysaker and Vestøl 2020). The initial point classification was done using the automated Terrain Classification plug-in module in CloudStation (YellowScan 2024a). This included classification of points originating from one of four classes: land (i.e. all points outside of the water), the water surface, water column noise, and sea floor (Supplementary Figure 2A). These point classes were also colourised in CloudStation using RGB data collected simultaneously by the Navigator’s onboard RGB camera (*n* = 218 images). Additional point attributes included scan angle, scan return number, and intensity. Raw intensity values were corrected during post-processing in CloudStation using the bathymetric LiDAR equation (Kashani et al., 2015; YellowScan, 2024a), and were reported on a scale between 0 and 5000.

### Data cleaning

While automatic point cloud classification in CloudStation expedited raw post-processing (Supplementary Figure 2A), further refinement was required to distinguish between true sea floor (including both sandy and rocky bottoms) and sea floor cover (i.e. vegetation) (Supplementary Figure 2B). Following raw post-processing, the point cloud was loaded into the open-source software CloudCompare (CloudCompare 2024) for inspection and further manual processing. We investigated classified sea floor and water column noise points and found that some points classified as water column noise in CloudStation originated from benthic vegetation, which was confirmed via visual inspection (see Supplementary Figure 2B). This was likely due to the elevation at which vegetation float within the water column relative to the sea floor. Filtering by intensity values (see next section), we retained points originating from vegetation and removed the remaining water column noise points manually, which were easily identified by visual inspection from the point cloud cross-sections (Supplementary Figure 2B). Final cleaning was also done in CloudCompare, where areas lacking sufficient sea floor point density were removed, based on the author’s discretion, to ensure reliable creation of digital models in subsequent analyses.

### Point cloud classification

In the present and subsequent sections, the code written for all analyses in R (R Core Team 2024) was assisted by the Claude Sonnet 3.5 Large Language Model (Anthropic 2024). To be able to create a dataset for classifying all points in a Random Forest model, we first manually annotate the point cloud by identifying vegetation and sea floor points based on intensity values from ground-truth observations. Manual inspection of the point cloud’s intensity values, at the location of ground-truth data points (Supplementary Figure 1), showed a marked difference between ‘true’ sea floor and vegetation intensity values (Supplementary Figure 2B). Guided by the ground-truth points, we extracted the range, mean, and standard deviation of eelgrass intensity values within the point cloud. We then annotated the point cloud data by labelling all points with intensity values less than the mean intensity value of eelgrass plus 1 standard deviation (S.D.) as vegetation, and points higher than this range as sea floor. This annotated dataset was used to train a random forest (RF) model for point classification.

Based hereon, we created a probability-based prior, with points within the pre-defined vegetation intensity range defined as > 90% probable of being vegetation, while points outside of this range as < 10% probable. In addition to the probability-based prior, we used R, G, and B values (each ranging from 0-255) derived from the simultaneously captured RGB imagery, as well as the nearest-neighbour distance (in cm) of points as predictor variables in the RF model to predict points as either sea floor or vegetation. The model was created using the randomForest package in R (Liaw & Wiener, 2002). The number of trees (ntree) was determined by the minimum Out of Bag (OOB) error across a range of 10 to 100 trees. The number of randomly drawn candidate variables was determined by grid search using the caret package (Kuhn, 2008). Data was split into 80% training and 20% testing subsets to evaluate final model performance via confusion matrix and precision scores. OOB error and variable importance (Mean Decrease Gini) were extracted using the randomForest package (Liaw & Wiener, 2002). After assessment, the resulting RF model was used to classify all points using the base R predict function (R Core Team 2024).

### Digital Terrain and Surface Models

Following point cloud classification, we created a DTM from the sea floor points, and a DSM from the vegetation points using the lidR R package (Roussel et al. 2020, Roussel & Auty, 2025). The sea floor DTM was created using the Inverse Distance Weighting (IDW) algorithm, using 10 nearest neighbour points and a weighting distance parameter of 2 (i.e. squared distance). The DTM was then smoothed in order to reduce the effects of fine-scale features of the bathymetry model that likely resulted from point cloud noise at the vegetation / sea floor interface (Wojciech 2022). Smoothing was completed using the focal function from the terra package in R (Hijmans 2024), applying a large moving window to ensure smoothness.

The vegetation DSM was created using the point-to-raster algorithm in the lidR package, which attributes the elevation of the highest point within each raster cell (Roussel et al. 2020, Roussel & Auty, 2025). As an objective of this study was to quantify eelgrass habitat volume, to prevent excessive voids within the modelled vegetation canopy DSM, we replaced each point with a ‘disk’ of 8 points surrounding the original point. This approach has been demonstrated to improve terrestrial forest canopy height models (CHMs) by removing ‘data pits,’ and as such the routine creates a DSM that captures the ‘outside’ of the canopy, thereby creating a smoothing effect on the DSM that align with the human perception of a canopy cover (Liu and Dong 2014). We assumed this would best align with traditional seagrass canopy height measures. To assess the sensitivity of this smoothing approach, we tested disk radii (i.e. smoothing window sizes) from 0.1 to 0.5 m at 0.1 m increments and evaluated their effects on estimated canopy height and volume.

Following the creation of digital models, we restricted the vegetation DSM to areas of eelgrass manually using ground-truth observations, in combination with visual assessment of the colourized LiDAR point cloud data and expert knowledge of the study area. Also, final DTM and DSM rasters were masked to overlapping extents between the 50 m and 25 m flight datasets, allowing for comparison. After restricting the DSM to areas of eelgrass, we calculated the cell-specific canopy height (CH_r_, in m) as the difference between the DSM and DTM elevations (z) of cell r:

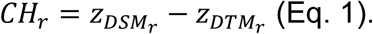

We then calculated the volume of eelgrass habitat across the study area (in m^3^) as the sum of the volume of all canopy cells (n), calculating the volume of cell r by multiplying canopy height (CH_r_, in m) by the area of each cell (m^2^):

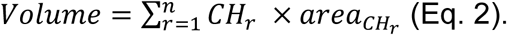

### Validation

We validated the LiDAR-derived elevation values for both terrestrial (above-water) and submerged (below-water) surfaces by comparing them with field measured ground-truth data. Validation metrics included mean error, standard deviation, and root mean square error (RMSE) between GNSS ground-truth data (handheld and echosounder) and LiDAR values. *In-situ* eelgrass canopy height was compared to LiDAR vegetation canopy height (i.e., the elevation difference between vegetation-classified LiDAR points and the DTM below the point). For all validations, we found the nearest LiDAR point to each ground-truth coordinates, based on Euclidean distance, using the nn2 function from the RANN package in R (Jefferis and others 2024).

### Eelgrass biomass and carbon content

We calculated eelgrass biomass and organic carbon content from our LiDAR-derived canopy model by incorporating the mean biomass per unit area calculated at the study site. Values were derived from eelgrass samples collected in September 2023, by manually measuring the height and wet weight of eelgrass in 20 × 20 cm cells (*n* = 42, Supplementary Table 1) (see Borger 2024). The biomass per unit area (hereafter referred to as volumetric biomass, B_vol_, g ww m^-3^) was then calculated using the equation:

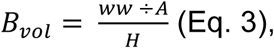

where ww equals the wet weight (g) of eelgrass samples, A equals the sample cell area (m^2^), and H equals the mean measured height of eelgrass (m) in the sample cell. We also report biomass g ww m^-2^ (ww/A). From the wet weight biomass values per unit area, we converted to dry weight (dw) using a conversion value derived from subsamples (*n* = 85, dw/ww = 0.24, Supplementary Table 2), and from dry weight to carbon (C/dw = 0.34) using standard C/dw conversion values from literature (Duarte 1990; Howard and others 2014). From *n* = 42 samples, we summarised the minimum, mean (S.D.), and maximum biomass per unit area values of eelgrass at the study area. In subsequent analysis, we assumed that this calculated range was representative of the eelgrass canopy at the time of LiDAR data sampling (August 2024).

We then calculated a range of total eelgrass biomass (ww, kg) across the study area using the following equation:

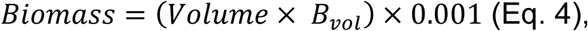

where volume equals the LiDAR-derived eelgrass habitat volume (m^3^), for both the 25 m and 50 m datasets. The range was calculated by applying the minimum, mean, and maximum B_vol_ values to Eq. 4. Using the conversion ratios, we converted total biomass ww (kg) to biomass dw (kg), and C (kg). For reporting purposes, we also summarised all biomass values m^-2^.

## Results

### LiDAR derived bathymetry DTM and DSM

For the 50 m LiDAR data, a total of 4,356,280 points were recorded across the 32,676 m^2^ study area following raw data post-processing (Figure 3A). After point cloud cleaning we retained 895,979 points across an area of 12,868 m^2^, with a mean density of 70 points per m^2^, 61 points per 1 m^3^ voxel (volume pixel), and a mean nearest neighbour distance of 8.9 cm (± <0.01 S.E.) (Figure 3B). From ground-truth data, points between 32 (minimum value) and 1263 intensity were annotated as vegetation points (eelgrass points mean intensity = 863 + 400 S.D., Figure 3C), with points above this range annotated as sea floor. The 25 m LiDAR dataset initially contained 5,682,638 points across 23,904 m^2^ (Figure 3D). After cleaning, 2,112,557 points remained across 16,840 m^2^ with a mean density of 125 points per m^2^, 105 points per 1 m^3^ voxel, and a mean nearest neighbour distance of 6.9 cm (± <0.01 S.E.) (Figure 3E). Points annotated as vegetation were those with intensity values from 100 (minimum value) to 1810 (eelgrass points mean intensity = 1512 + 298 S.D., Figure 3F), and points above this range were annotated as sea floor. The depth range of both cleaned datasets extended from 0 cm (i.e. the land-water interface) down to approximately 3 m (Supplementary Figure 1).

**Figure 3:**
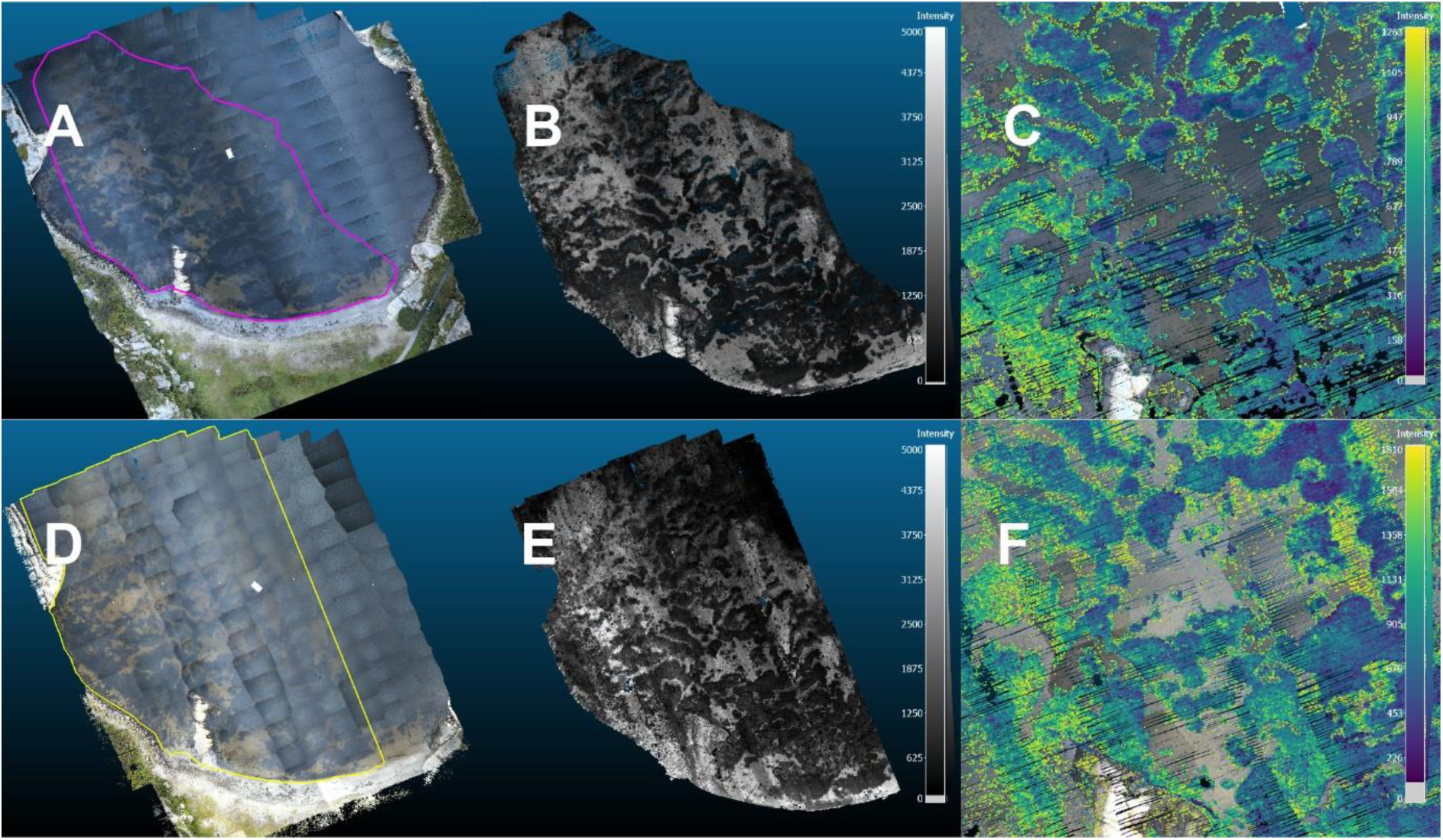
A) A 2D representation of the full point cloud from the 50 m LiDAR dataset, comprising 4,356,280 individual points accurately positioned in 3D (x, y, z). Points are colourized from concurrently collected RGB images (n = 218). The pink outline indicates the subsection shown in B. B) The cleaned 50 m point cloud (n = 895,979) used to create a 3D representation of the eelgrass (*Zostera marina*) meadow and calculate canopy height, habitat volume, and biomass. Points are scaled by their intensity values between 0 – 5000. C) The intensity scale of the 50 m, annotated vegetation points before classification (colour-scaled points, intensity ≤ 1263). D) The full 25 m LiDAR dataset, comprising 5,682,638 points, with the yellow outline indicating the subsection in E. E) The cleaned 25 m point cloud (n = 2,112,557), scaled by intensity values. F) The intensity scale of the 25 m, annotated vegetation points before classification (colour-scaled points, intensity ≤ 1810). This figure was created using CloudCompare (2024).

From the resulting point clouds, the RF models used for point cloud classification showed 100% accuracy in distinguishing between sea floor and vegetation within our test datasets. Both the 50 m and 25 m models showed low OOB error and classification error rates, converging to 0 before reaching 40 trees (Supplementary Figure 3A and 3B). The intensity-based probability-prior was by far the most important variable in predicting point classification (Supplementary Figure 3C and 3D). The final RF models were then used to classify all points as either sea floor or vegetation, allowing generation of the DTMs and DSMs.

The DTMs and DSMs from both 50 m and 25 m LiDAR datasets were created at a resolution of 10 cm, which is slightly coarser than the average point spacing of 8.9 cm in the 50 m dataset. The vegetation DSMs were restricted to areas of eelgrass, removing canopy coverage of macroalgae species (primarily *Fucus serratus* and *Fucus vesiculosus*) growing on hard substrate surrounding a bedrock outcropping in the southwestern corner of the bay (Figure 3 and 4). Subtracting the DTM from the DSM resulted in the eelgrass canopy height model (Figure 4).

**Figure 4:**
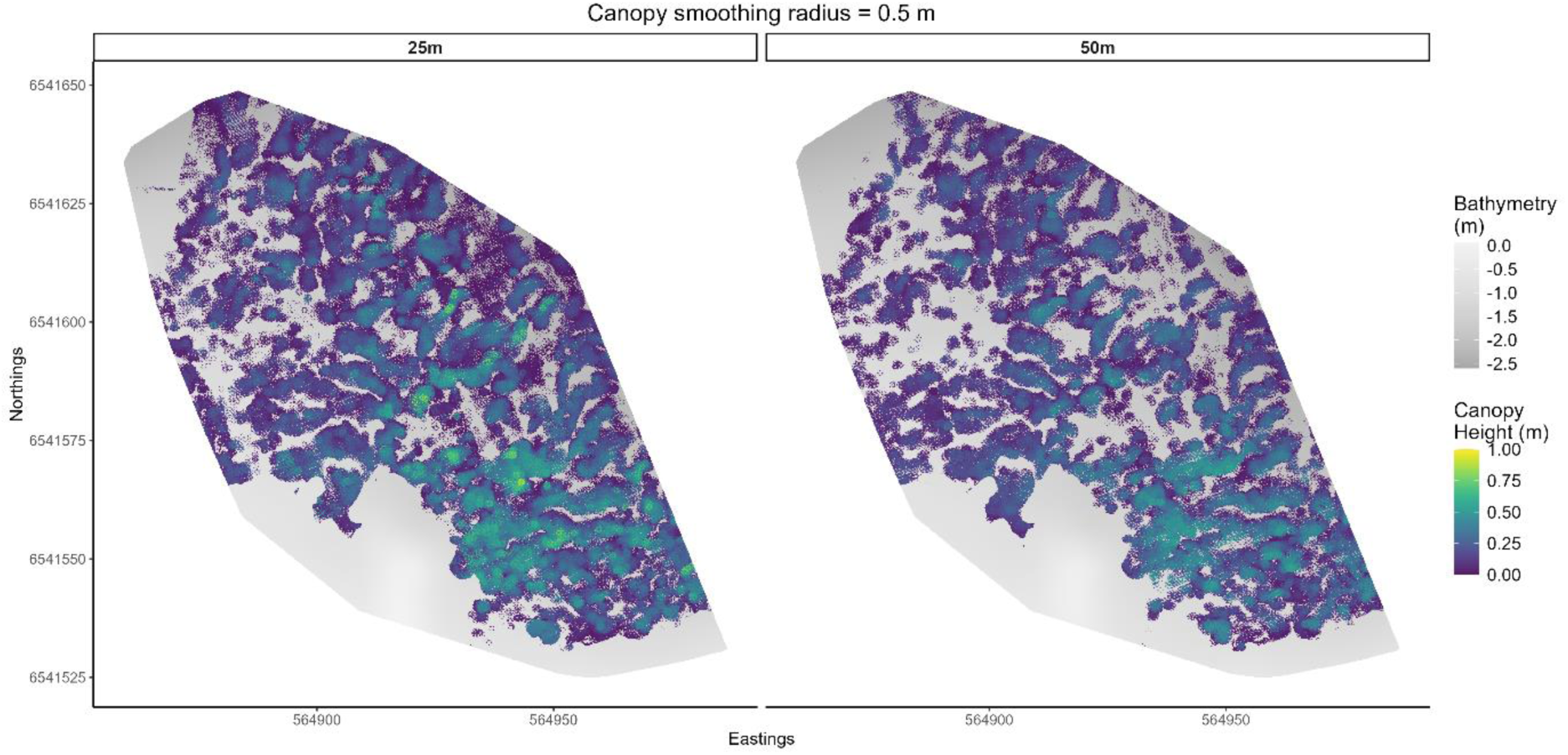
The Digital Surface Model, coloured from purple (low) to yellow (high), representing the eelgrass (*Zostera marina*) canopy, overlaying the Digital Terrain Model (in grey), which shows the sea floor bathymetry (m) in Ølbergholmen, Norway, for the 25 m and 50 m LiDAR datasets. Canopy height (m) is calculated as the difference between the DSM and DTM. Digital models were created using the lidR package in R (Roussel et al. 2020, Roussel & Auty, 2025), with a canopy smoothing radius = 0.5 m.

Comparing LiDAR-derived sea floor elevations with *in-situ* GNSS measurements demonstrated a strong consistency between the 50 m LiDAR dataset and ground truth elevation, showing a mean vertical error of only 0.5 cm (12.0 cm S.D., 11.7 cm RMSE) (Table 1, Figure 5). The 25 m dataset showed a larger offset than the 50 m dataset (mean vertical error of 6.5 cm, 15.4 cm S.D., 16.3 cm RMSE) (Table 1, Figure 5). When compared with echosounder-derived sea floor bathymetry, the 25 m LiDAR dataset showed better agreement than the 50 m dataset, with a 10 cm mean vertical error (8.9 cm S.D., 13.4 cm RMSE) (Table 1, Figure 5). For terrestrial LiDAR points (i.e. on land outside of the study area), both datasets produced similar vertical accuracy (mean error = 0.6 cm), with the 25 m dataset displaying slightly lower S.D. (3.5 cm) and RMSE (3.5 cm) (Table 1, Supplementary Figure 4).

**Figure 5:**
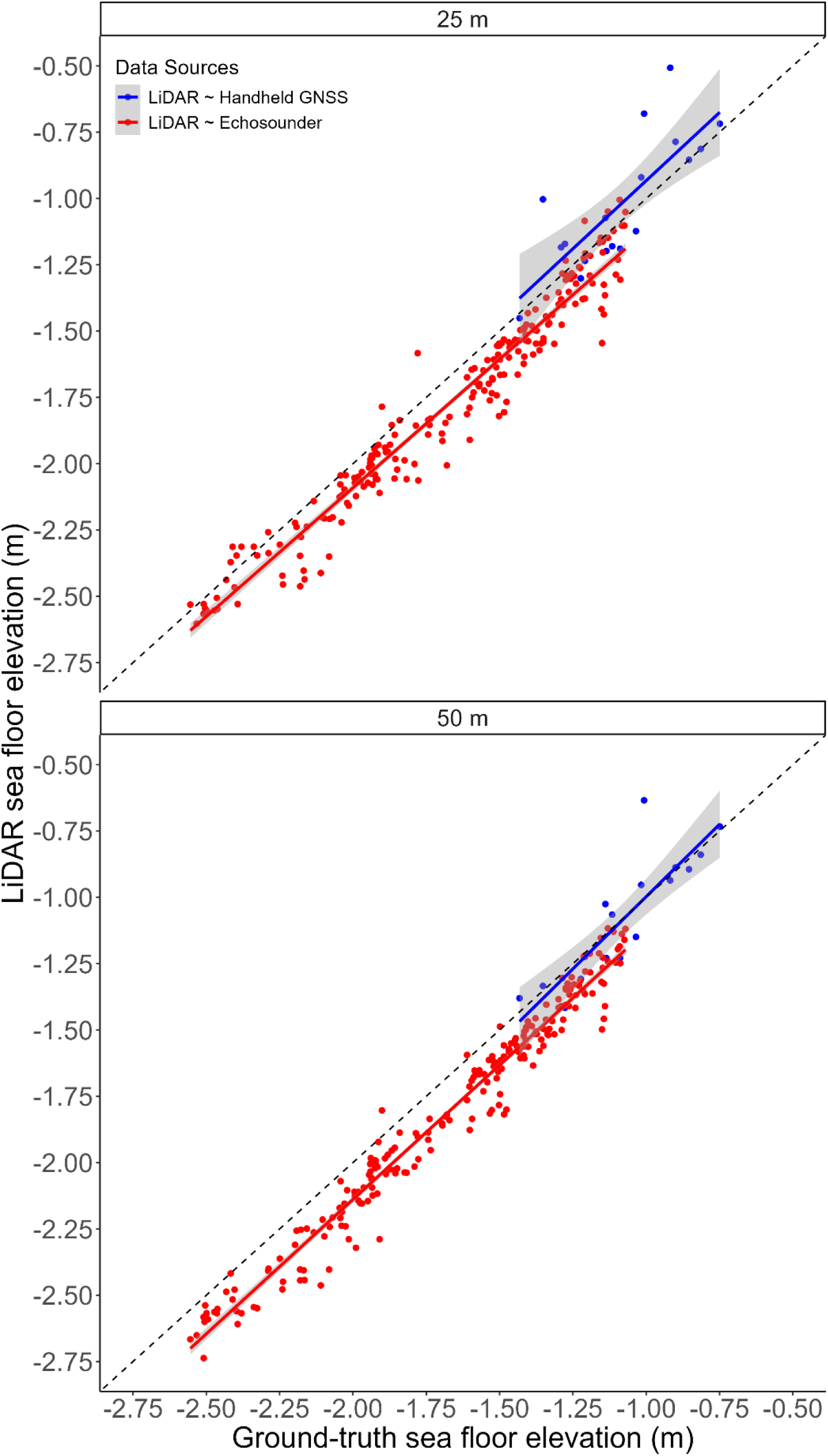
Correlation between the LiDAR-derived bathymetry and ground-truth data (m) collected from handheld positioning Global Navigation Satellite System (GNSS) receiver (blue) and echosounder (red) for A) the 25 m and B) the 50 m LiDAR flight elevation datasets. The dotted line represents a 1:1 relationship.

**Table 1:**
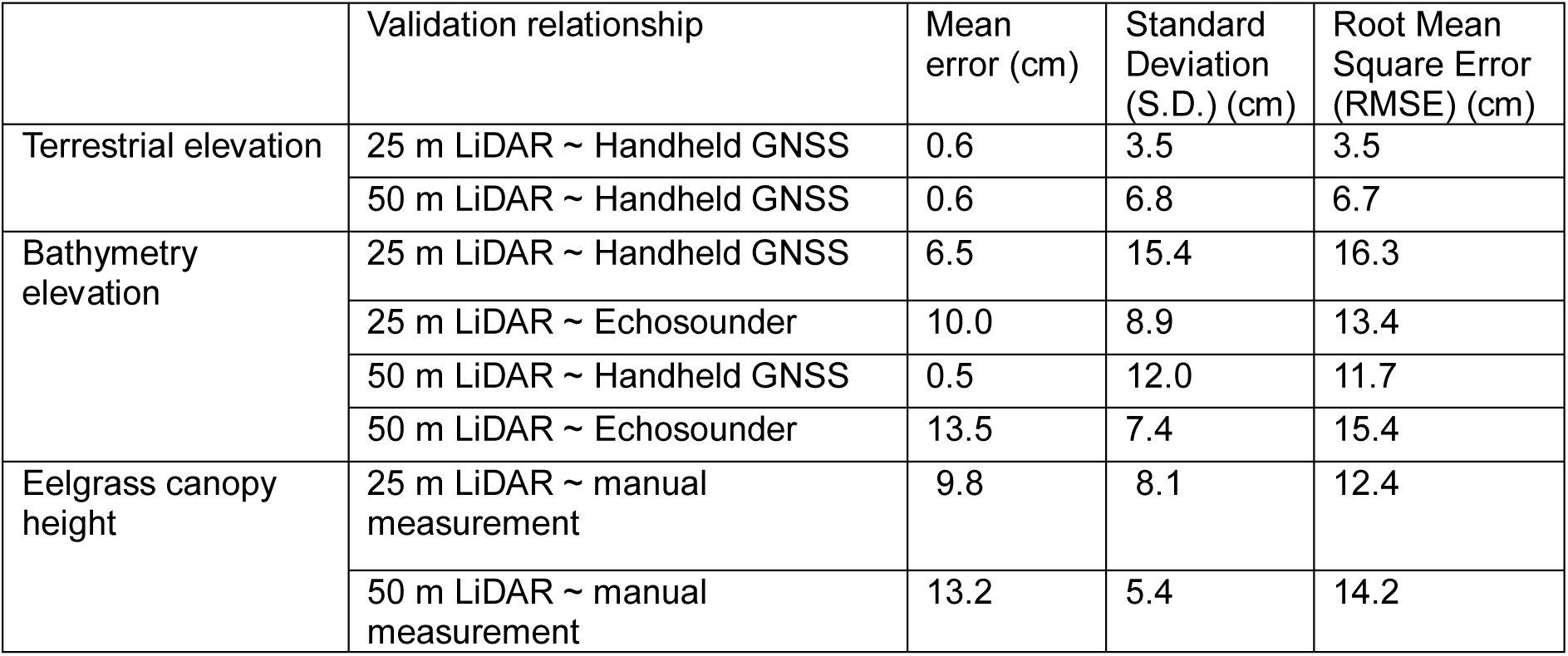
Validation of LiDAR elevation. LiDAR-derived terrestrial elevation, bathymetry elevation, and eelgrass canopy height, relative to ground-truth values. Eelgrass canopy height was compared to manual measurements, bathymetry was compared to handheld Global Navigation Satellite System (GNSS) elevation measurements and echosounder derived bathymetry, while terrestrial elevation (outside of the focal study area) was compared with handheld GNSS measurements only. Validation is reported as the mean vertical error, standard deviation (S.D.) and Root Mean Square Error (RMSE) in centimetres between LiDAR elevation values and ground-truth values.

### Seagrass canopy height and meadow volume

Canopy heights derived from both the 50 m and 25 m LiDAR datasets (from late August 2024) were generally lower than manually measured ground-truth data (from early November 2024) (Figure 6A). Comparing the eelgrass canopy heights by manual measurements between November 2024 and September 2023, canopy heights were slightly higher in September 2023 than November 2024, with a strong positive relationship between the two measurements, baring one outlier (Supplementary Figure 5).

**Figure 6:**
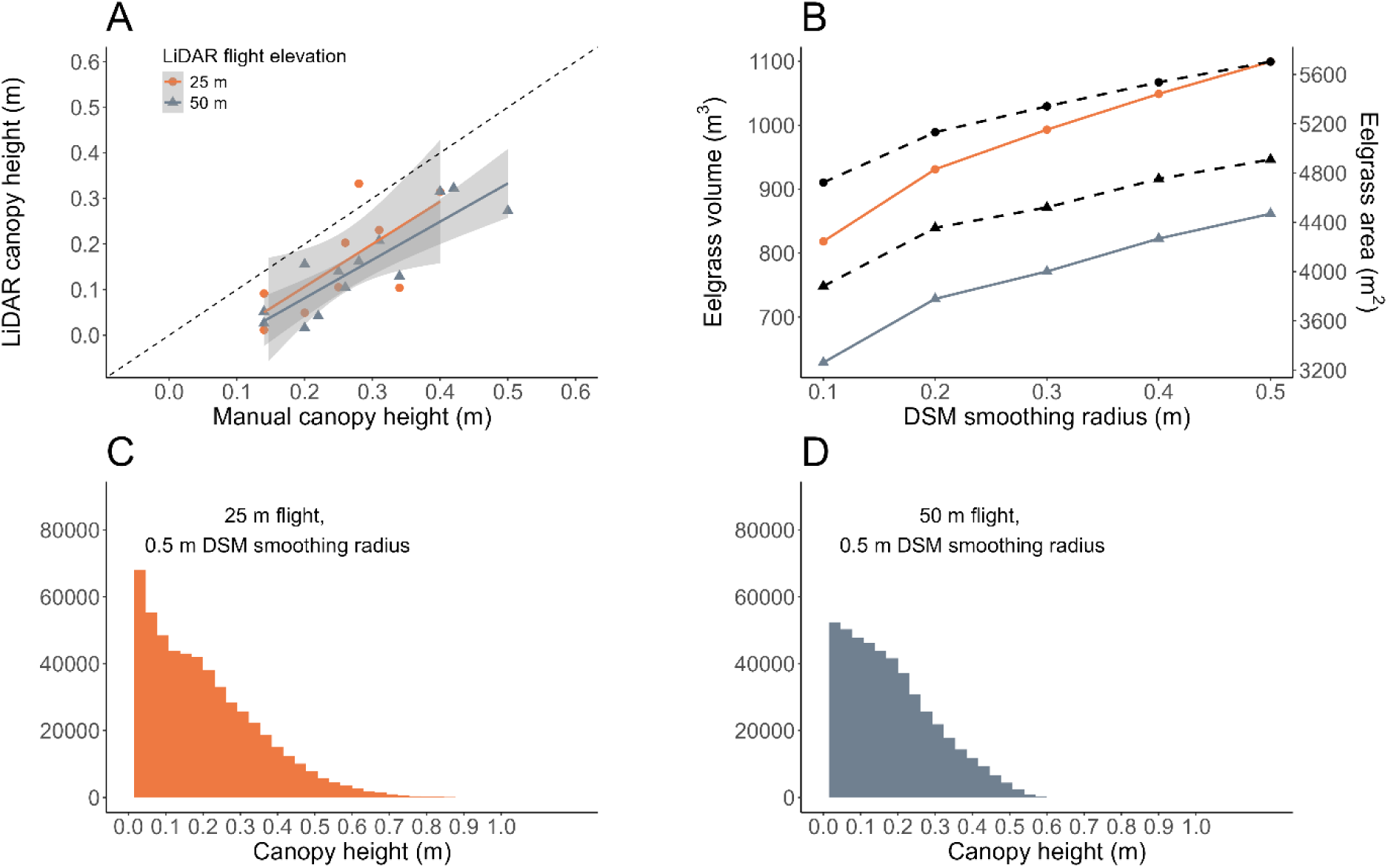
A) LiDAR-derived eelgrass canopy height (m), acquired in August 2024 at 25 m and 50 m flight elevation, plotted against manual canopy height measured *in-situ* (m) collected in November 2024. The dotted line represents a 1:1 relationship. B) Total eelgrass habitat volume (solid-coloured lines, left, m3) calculated from the 25 m and 50 m elevation datasets, plotted against Digital Surface Model (DSM) smoothing radius (m). Also plotted is the total eelgrass area (dashed black lines, right, m2) for both LiDAR datasets. Plotted are histograms of canopy height (m) raster cells for both the C) 25 m and the D) 50 m LiDAR datasets. For both histograms, results are from DSMs with a 0.5 m smoothing radius.

Modelled eelgrass canopy height, area, and volume were influenced by both flight elevation and the smoothing radius of the DSM (Figure 6A and 6B). Total eelgrass canopy area and volume were higher for the 25 m datasets, while increasing the DSM smoothing window increased canopy area and volume for both datasets (Figure 6B). Canopy area ranged from 4723 to 5703 m^2^ (mean = 5287, S.D. = 381) and volume from 818 to 1099 m^3^ (mean = 978, S.D. = 109) across smoothing radii for the 25 m dataset, while area ranged from 3881 to 4909 m^2^ (mean = 4484, S.D. = 398) and volume from 629 to 862 m^3^ (mean = 763, S.D. = 90) for the 50 m dataset (Figure 6B). With the largest smoothing radius (0.5 m), canopy height ranged from 0.0 to 0.98 m for the 25 m dataset (Figure 6C), and from 0.0 to 0.64 m for the 50 m dataset (Figure 6D). The distribution of canopy height cells displayed a general increase concurrent with an increase in smoothing radius (Supplementary Figure 6). When comparing canopy height and bathymetry elevation between the 50 m and 25 m datasets (Supplementary Figure 7A), the 50 m dataset is generally lower (Supplementary Figure 7B). Moreover, the areal coverage of eelgrass canopy height raster cells is higher within the 25 m dataset (Supplementary Figure 8).

### Above ground eelgrass biomass and carbon

*In-situ* eelgrass volumetric biomass ranged from 84.5 to 3446.6 g ww m^-3^ with a mean of 1363.5 g ww m^-3^ (S.D. = 999.1) (Table 2, Supplementary Table 1). Using these values we then calculated the range of total biomass values (total kg ww, kg ww m^-2^) across the study area for both the 25 m and 50 m datasets (Table 3). After converting to dry weight, then carbon content, we estimate a range between 7.6 kg to 309.2 kg, with a mean of 122.3 kg C content from the 25 m dataset, and a range between 5.9 kg and 242.3 kg, with a mean of 95.9 kg C content from the 50 m dataset (Table 3). This equates to approximately 20 g C m^-2^ (50 m dataset) or 21 g C m^-2^ (25 m dataset) (Table 3).

**Table 2:** Eelgrass biomass per unit area. Results deriving the mean (and Standard Deviation, S.D.), minimum and maximum biomass per unit area from samples collected at the study site (*n* = 42) in September 2023 (see Supplementary Table 1). This includes the biomass density in grams (g) of wet weight (ww) m^-2^ and m^-3^, the biomass density in g dry weight (dw) m^-2^ and m^-3^, and biomass in g carbon (C) m^-2^ and m^-3^. Wet weight to dry weight ratios were derived from subsamples (dw/ww = 0.24, Supplementary Table 2), while a standard dry weight to carbon ratio was used from literature (C/dw = 0.34).

| | Biomass (g ww $\text{m}^{-2}$) | Biomass density (g ww $\text{m}^{-3}$) | Biomass (g dw $\text{m}^{-2}$) | Biomass density (g dw $\text{m}^{-3}$) | Biomass (g C $\text{m}^{-2}$) | C density (g C $\text{m}^{-3}$) |
| --- | --- | --- | --- | --- | --- | --- |
| Mean (S.D.) | 364.1 (343.7) | 1363.5 (999.1) | 86.3 (81.5) | 323.1 (236.8) | 27.6 (26.1) | 103.4 (75.8) |
| Minimum | 19.4 | 84.5 | 4.6 | 20.0 | 1.5 | 6.4 |
| Maximum | 1353.0 | 3446.6 | 320.7 | 816.8 | 102.6 | 261.4 |

**Table 3:**
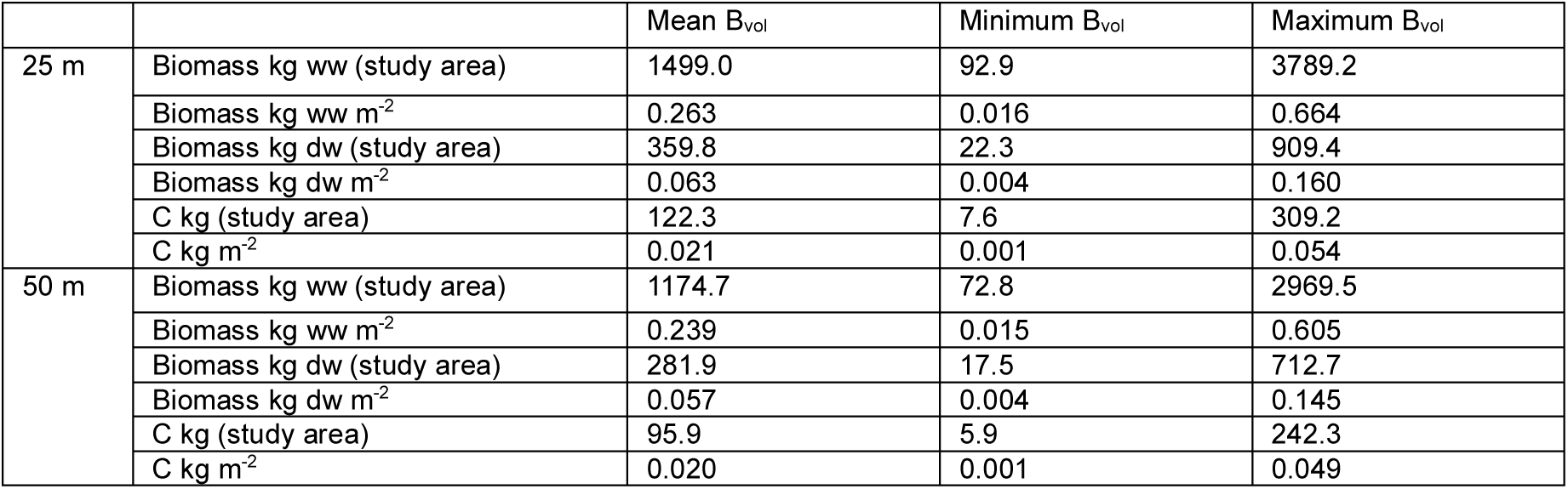
Summary of eelgrass biomass across the study area. Results summarising the range of eelgrass biomass across the study area, by applying the mean, minimum and maximum volumetric biomass (B_vol_) values, to both the 25 m and 50 m LiDAR-derived eelgrass canopy model. Included is the total biomass (kg) in wet weight (ww) across the study area, the biomass (kg) ww m^-2^, biomass dry weight (dw) across the study area, biomass dw m^-2^, biomass (kg) of carbon (C) across the study area, and biomass (kg) C m^-2^. Results displayed here are from canopy height models with a DSM smoothing radius of 0.5 m only.

## Discussion

From drone-borne topobathymetric LiDAR data, we were able to estimate the areal extent and the specific canopy height of an eelgrass meadow habitat within a confined coastal study area at a spatial resolution of ∼10 cm. By creating high-resolution digital models of both the sea floor and the overlaying eelgrass canopy, the habitat volume, biomass and above-ground carbon stock can be estimated. These novel results build upon previous applications of airborne topobathymetric LiDAR for mapping the spatial extent of marine vegetation and classifying habitat types (Pan and others 2014; Ishiguro and others 2016; Letard and others 2021, 2022; Ekelund and others 2024). However, this study advances these applications by deriving and analysing the three-dimensional geometric structure of submerged vegetation, which enable quantification of habitat volume and seagrass biomass to provide remotely sensed estimates of blue carbon stocks.

Airborne LiDAR has been applied extensively in terrestrial forest research (Coops and others 2021). For example, LiDAR-derived forest canopy structure has been related to shading and microclimatic conditions – factors that directly influence plant growth, competition, and community composition (Zambrano and others 2019; Zellweger and others 2019). Airborne LiDAR also provides accurate tree height and crown area measurements at the scale of entire forests, facilitating large-scale assessments of biomass and carbon storage (Lines and others 2022). Our results demonstrate the potential of topobathymetric LiDAR to facilitate similar ecological investigations in shallow coastal waters, enabling structural analyses and carbon stock estimations for submerged vegetation such as seagrass.

Estimation of living biomass and blue carbon stocks of above-ground seagrass meadows in cost-efficient manners are an inherent challenge to monitoring and mapping actions (Boström and others 2014). Traditionally it is performed by labour intensive methods, and sufficient data is often a limiting factor to nature protection agendas and ecosystem accounting programs (Carruthers and others 2024). LiDAR point cloud data advances quantitative estimation of biomass and blue carbon from remotely sensed data as a cost-efficient tool, as we here demonstrate showing a method for digital, systematic, and remotely sensed assessment. The biomass of the entire eelgrass meadow was calculated considering both a precise estimate of the areal extent and the canopy height at a spatial resolution of ∼10 cm. This is in large contrast to traditional measurements where spatial distribution is often based on point measurements of seagrass presence/absence along predefined transects, with canopy height determined from manual measurements using SCUBA or snorkelling at random locations inside the meadow (Neckles and others 2012).

Still the estimate holds uncertainties, including a known volumetric biomass density, and an accurate LiDAR derived canopy height. The biomass density of the canopy is influenced by the shoot density, leaf length and weight of individual plants (Boström and others 2014). Here we estimated the biomass density from the weight and height of the canopy at several locations in the study area (*n* = 42), and thus assumed an average relationship between leaf length, shoot density, and weight. The shoot density and above-ground biomass have shown to be correlated, however showing large variabilities, while per meter biomass becomes independent of shoot density as it approaches maximum biomass (Boström and others 2014). Largely, biomass and cover has shown to depend on light availability and thus depth (Sand-Jensen and Borum 1991; Krause-Jensen and others 2000). In the present study all data were sampled at depths shallower than 3 meters. Our sampled biomass density had a mean of 323.1 g dw m^-3^ with a standard deviation of 236.8 g dw m^-3^ (Table 2). As such, the uncertainty of the biomass estimate should be considered, whilst the accuracy of the LiDAR derived canopy height is discussed further below. We recommend that the relationship between *in-situ* eelgrass biomass density and drone-borne LiDAR point cloud density be investigated in further studies.

Eelgrass biomass in the Nordic countries exhibits considerable spatial and seasonal variability, influenced by factors such as depth, light availability, and nutrient conditions (Boström and others 2014; Krause-Jensen and others 2022). Our estimated range of above-ground biomass (4 to 160 g dw m^-2^, Table 3) overlaps within a previously reported range of 26 to 546 g dw m^-2^, also close to values reported in September in the area (∼ 170 g dw m^-2^) (Carstensen and others 2016). This variability reflects the dynamic nature of these coastal ecosystems and underscores the importance of site-specific data for accurate carbon budgets and management actions. Overall, the mean above ground carbon stock was estimated between 95.9 kg C (50 m) and 122.3 kg C (25 m) at the study site (or between 20 and 21 g C m^-2^, Table 3), corresponding to 352.0 to 448.8 kg CO2 stored in a section of the small bay.

Biogenic habitats such as seagrass meadows play an important role in supporting biodiversity in coastal ecosystems (McHenry and others 2021; Duarte and others 2025). Seagrasses modulate hydrodynamic processes by influencing currents, wave propagation, and erosion rates (James and others 2020; van de Vijsel and others 2023), thereby providing hydrodynamic shelter for a large number of species living epiphytically on the plants or within the plant canopy (Castejón-Silvo and others 2021; Barcelona and others 2024). Most studies on biodiversity in seagrass habitats have compared vegetated and unvegetated areas (Rodil and others 2021) or focused on the effects of meadow size, patchiness, and edge effects (Boström and others 2006; Duffy 2006). Estimating the biomass and canopy height of seagrass meadows has been more challenging, and there are concurrently fewer studies investigating the role of the three-dimensional structure of the habitat for the fauna community. Nevertheless, both faunal abundance and biodiversity have been shown to correlate positively with seagrass biomass and canopy height (Orth and others 1984; Rodil and others 2021).

Our results demonstrate the potential of drone-borne LiDAR to provide detailed, three-dimensional structured data on submerged vegetation, that can be related to associated fauna. We recommend for further research to combine LiDAR-derived habitat structure with automated biodiversity monitoring techniques, such as remotely operated vehicles (ROVs) and machine learning (Goodwin and others 2022; Knausgård and others 2022; Vigo and others 2023), to promote a fully automated and reproducible ecological monitoring of marine biodiversity.

Our results provide a range of estimated eelgrass habitat volumes within the analysed area (Figure 6B). By comparing LiDAR-derived canopy heights with ground-truth measurements, we found that *in-situ* values were generally higher (Figure 6A). This pattern is in line with previous studies in terrestrial grasslands, whereby LiDAR-derived canopy height and estimated aboveground biomass tended to underestimate field observations (Miura and others 2019; da Costa and others 2021). Reduced LiDAR canopy height estimates may result either from insufficient point returns at the canopy top (Miura and others 2019), likely due to the small surface of grass blades being missed by the LiDAR system (Zhao and others 2022), or from a lack of ground return points at the bottom, as dense canopy cover prevents laser penetration down to the ground (Hu and others 2020). Since our LiDAR data were collected around the seasonal peak of aboveground eelgrass biomass in the region, the raw point cloud revealed little to no sea floor return points under thick patches of eelgrass (see Supplementary Figure 2). Such dense canopies can therefore create gaps of terrain points, influencing the accuracy of the DTM interpolation (Cățeanu & Ciubotaru, 2021; Moudrý et al., 2020) and thereby influencing calculated canopy height.

Differences in modelled canopy height and sea floor bathymetry between the 50 m and 25 m LiDAR datasets further highlight the influence of canopy loss from both canopy top and/or bottom. With nearly double the point density per m2, the 25 m LiDAR dataset likely captured both more returns at the canopy top, as well as sea floor between eelgrass plants (Getzin and others 2022), thus explaining the variation in modelled canopy extent, height and DTMs between the two LiDAR datasets (Supplementary Figures 6-8). Moreover, variation in DTMs could also result from more noise present within the higher-density 25 m point cloud data (Liao and others 2024). These results, along with the influence of the DSM smoothing window, highlight the influence of UAV flight elevation, point cloud density, and digital model parameters in determining geometric habitat volume from drone-borne LiDAR, and thus, should be considered in future application.

Validation of LiDAR elevation values using echosounder bathymetry revealed a vertical offset of approximately 14 cm (RMSE, Table 1), although the linear relationship between the two methods was strong (Figure 5). This offset may reflect methodological differences in how elevation is calculated between LiDAR and sonar, for example, data registration, noise filtering, terrain characterisation, and the distance between the sensors and the sea floor (Li and others 2022; Raj and others 2024). Both LiDAR and sonar have their own advantages and limitations. Sonar can acquire bathymetric data at greater depths and is less affected by water quality, but its accuracy will be limited in shallow waters with complex terrain (Dudkov and Dorokhova 2020). Conversely, while drone-borne LiDAR cannot obtain data in deeper waters (as was the case in deeper sections of our study area), it can effectively survey the land-water interface and complex terrain, cover a larger geographic area, as well as derive radiometric information of surveyed surfaces. In our results, we demonstrate the benefits of drone-borne topobathymetric LiDAR as an efficient, high-resolution, and cost-effective way to survey shallow coastal areas and submerged vegetation, which are critical habitats for biodiversity and blue carbon storage.

Furthermore, our use of a commercially available, medium-sized drone with an integrated topobathymetric LiDAR system presents benefits relative to traditional Airborne LiDAR Bathymetry (ALB) on crewed aircraft surveys. ALB surveys usually rely on sensors fixed to crewed airplanes, which are associated with higher operating costs and reduced reproducibility (i.e. not pre-programmable) compared to drone deployment. Although ALB surveys currently cover larger areas, it generally produces lower point cloud densities. For example, Ekelund et al. (2024) surveyed seagrass beds across ∼1100 km^2^ via ALB, achieving an average point cloud density of 4 sea floor points per m^2^. While no doubt effective for surveying large areas, ALB resolution may limit structural analysis. In contrast, our drone-borne LiDAR survey, covering a much smaller area, produced an average point density of 70 points per m^2^ of sea floor (50 m dataset), facilitating detailed, high-resolution investigation of marine vegetation structure. Flying the drone at a higher elevation (120 m maximum height, according to the manufacturer), could provide a wider areal coverage, but reduced point density and thus variation in geometric estimates from the point cloud. Our results display the importance of such considerations when mission planning to monitor submerged aquatic vegetation.

In the present study, point cloud classification was limited to two classes (sea floor and vegetation), with high accuracy classification between the two. Guided by annotations informed via ground-truth data, these results were to be expected, as sandy / rocky bottom will reflect more light than plant surfaces, which tend to absorb more light (Garroway and others 2011). Following classification and subsequent digital model creation (DTM and DSM), we manually removed areas containing other types of vegetated habitats. To circumvent this and leverage a computational workflow, we recommend that future applications include multiple vegetation types, based on ground-truth radiometric values specific to individual species or species groups. As well, habitat classifications derived from drone-borne photogrammetry and machine learning (ML) can be used to guide point cloud classification or digital model segmentation (Gundersen and others 2024; Kvile and others 2024).

We consider that green LiDAR holds the potential to quantify the lower depth limit of seagrass growth and its relation to bathymetry and environmental gradients, a key indicator of ecosystem status within the European Commission’s Marine Strategy Framework Directive (European Commission 2008; Natural England 2023). We did not explore the depth penetration relative to light attenuation in this study. However, the green LiDAR instrument used here is consider capable of reaching a depth of ∼2 Secchi depth, which is within the range of reported lower growth limit for eelgrass; typically, at 1 to 3 Secchi depth (Nejrup and Pedersen 2008) This however remains to be empirically tested and Secchi depth only represents an approximation of light attenuation. (Krause-Jensen and others 2011). Drone-borne topobathymetric LiDAR also displays the potential to further map other important submerged habitats and blue carbon ecosystems, including coral reefs (Harris and others 2023) mussel or oyster reefs (Espriella and others 2023), maerl beds and kelp forests. As well, coastal erosion studies, sediment relocation and changes in bathymetry and ecosystem structure are areas to which topobathymetric LiDAR can improve and contribute to through further development for future applications.

The potential benefits of drone-borne green LiDAR in environmental and ecological management of shallow coastal habitats are substantial. Our results display the ability of this technology to map areas at high resolution (cm-scale), to distinguish between hard bottom and vegetative habitat, and to create 3D models of geometric structure. From these models, we were able to estimate habitat volume and biomass of above- ground living eelgrass tissue, thus facilitating blue carbon content assessments. Considering eelgrass’ shallow water habitat and patchy distribution, we display the utility and cost-effectiveness of drone-borne LiDAR relative to more traditional methods, such as manual *in-situ* monitoring, echosounders, or ALB surveys. This utility also has the potential to advance ecological research and inform management decisions, for example by providing key parameters, such as canopy height and volume and extent.

## Data availability

Code for the present analysis is available at the corresponding author’s GitHub (https://github.com/charles-patrick-lavin/NIVA-SeaBee-LiDAR), while the data analysed are available upon request.

## Acknowledgements

This work was funded by the Research Council of Norway and is a product of SeaBee (Norwegian Infrastructure for drone- based research, mapping and monitoring in the coastal zone, RCN project ID #296478). Additional funding was received from HORIZON-CL5-2023-D1-02 (C-BLUES) and HORIZON-CL6-2022-BIODIV-01-01, Grant agreement #101081642 (OBAMA-NEXT). We would also like to thank Christian Lindermann for assistance during field sampling.

---

**Supplementary Figure 1:**
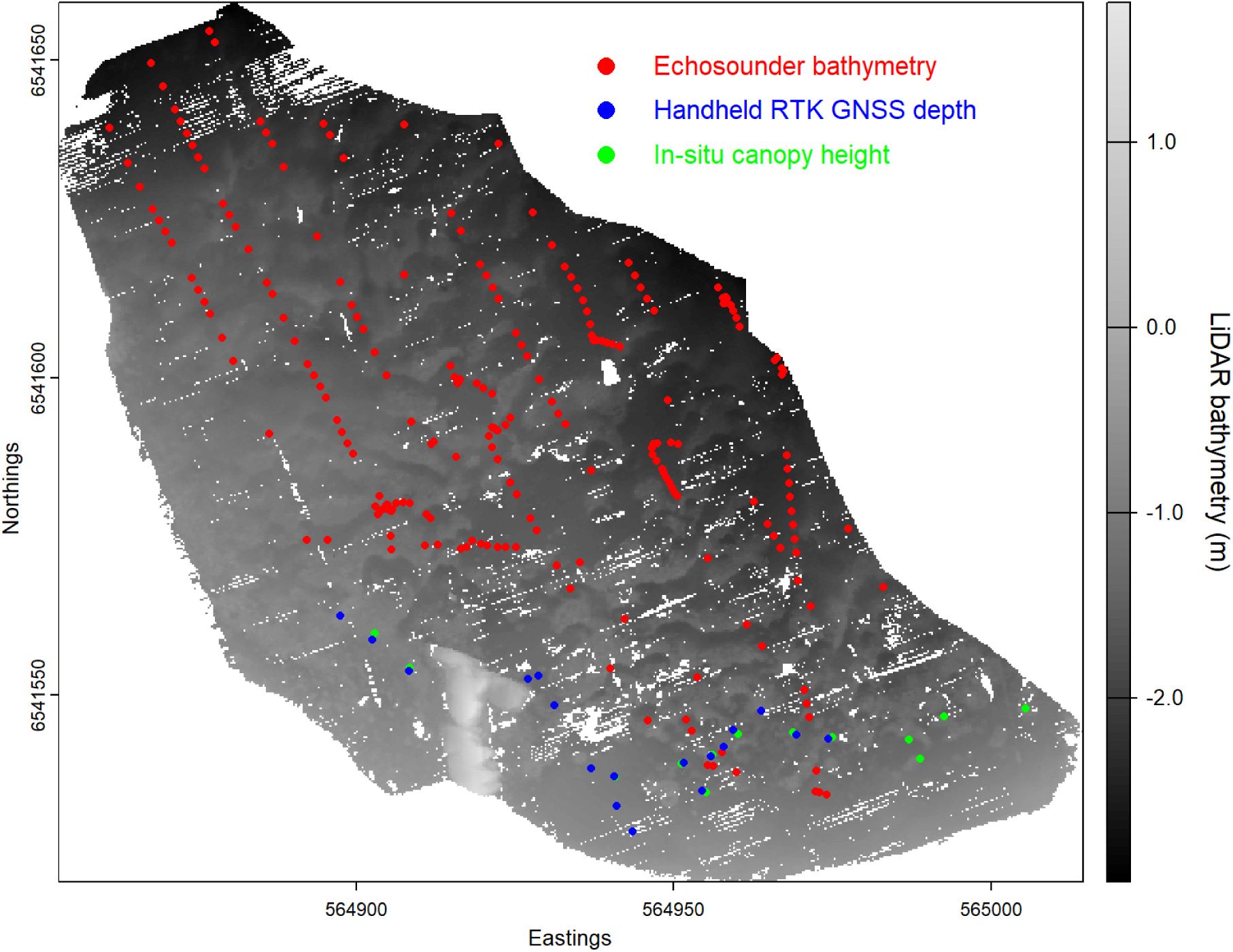
LiDAR-derived sea floor point cloud (50 m dataset) of the study area at Ølbergholmen, colorised by bathymetric depth, ranging from deepest (∼-3 m, dark grey) to shallowest (1.5 m, light grey). The following ground-truth validation points are overlaid: echosounder bathymetry (red), handheld RTK GNSS depth measurements (blue), and *in-situ* eelgrass canopy height measurements (green).

**Supplementary Figure 2:**
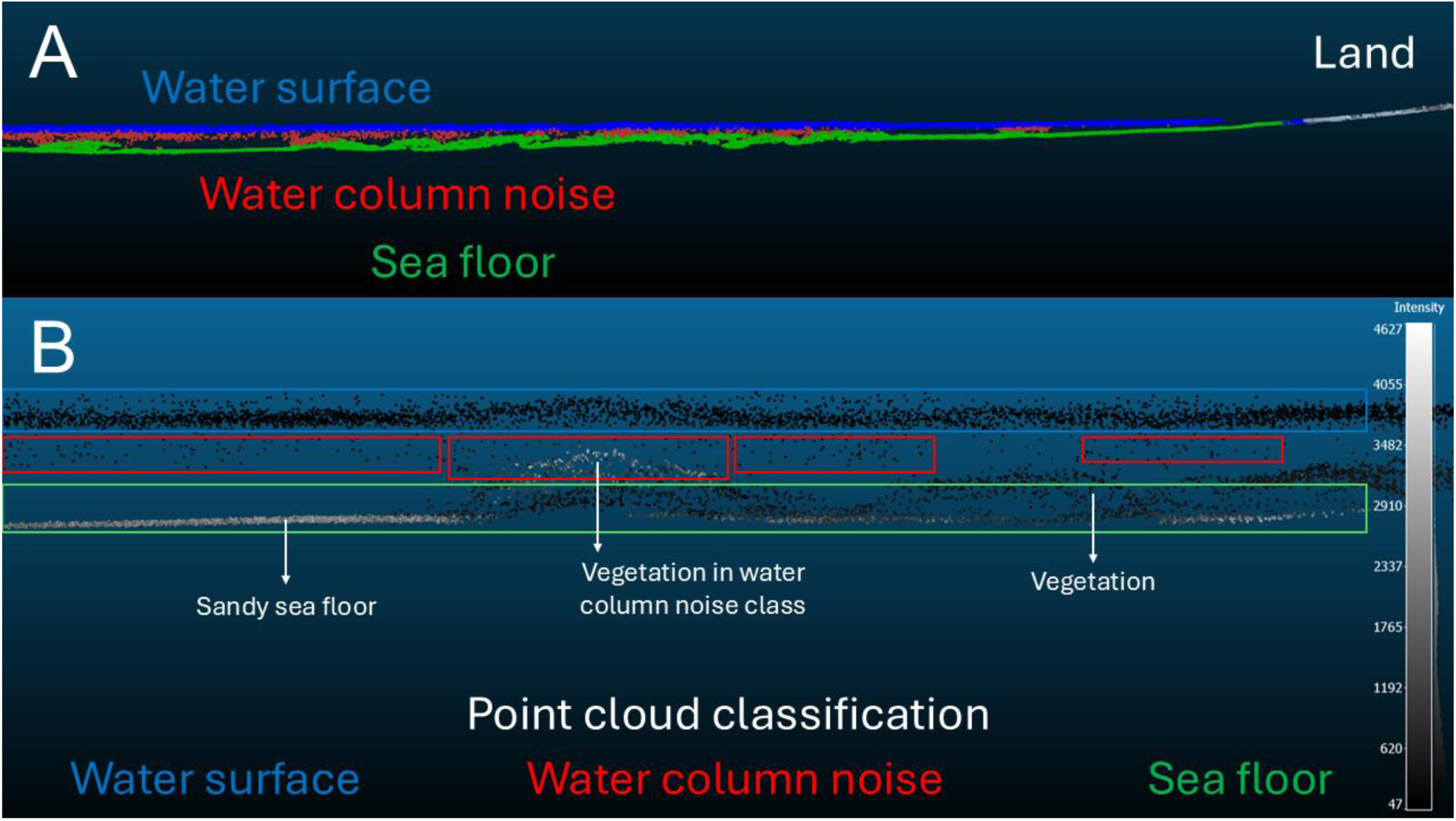
A cross-section of the LiDAR data, displaying A) the point cloud classifications generated using the YellowScan CloudStation software, including: land (white), the water surface (blue), water column noise (red), and sea floor (green). B) A close-up of Figure A highlighting the different classes including the water surface (blue) and water column noise (red), which included ‘true’ noise points, as well as some vegetation points classified as water column noise. After identifying vegetation points contained in water column noise, these points were included in the later analysis as vegetation points, while the rest of noise points were removed. The sea floor classification (green) contained sea floor cover points (e.g., vegetation), as well as ‘true’ sea floor point (e.g., sandy sea floor). Classification between true sea floor and vegetation points was completed via random forest (RF), including intensity values, R, G, and B values (colourised from the drone’s onboard RGB camera), as well as nearest neighbour distance (cm) from points as model variables.

**Supplementary Figure 3:**
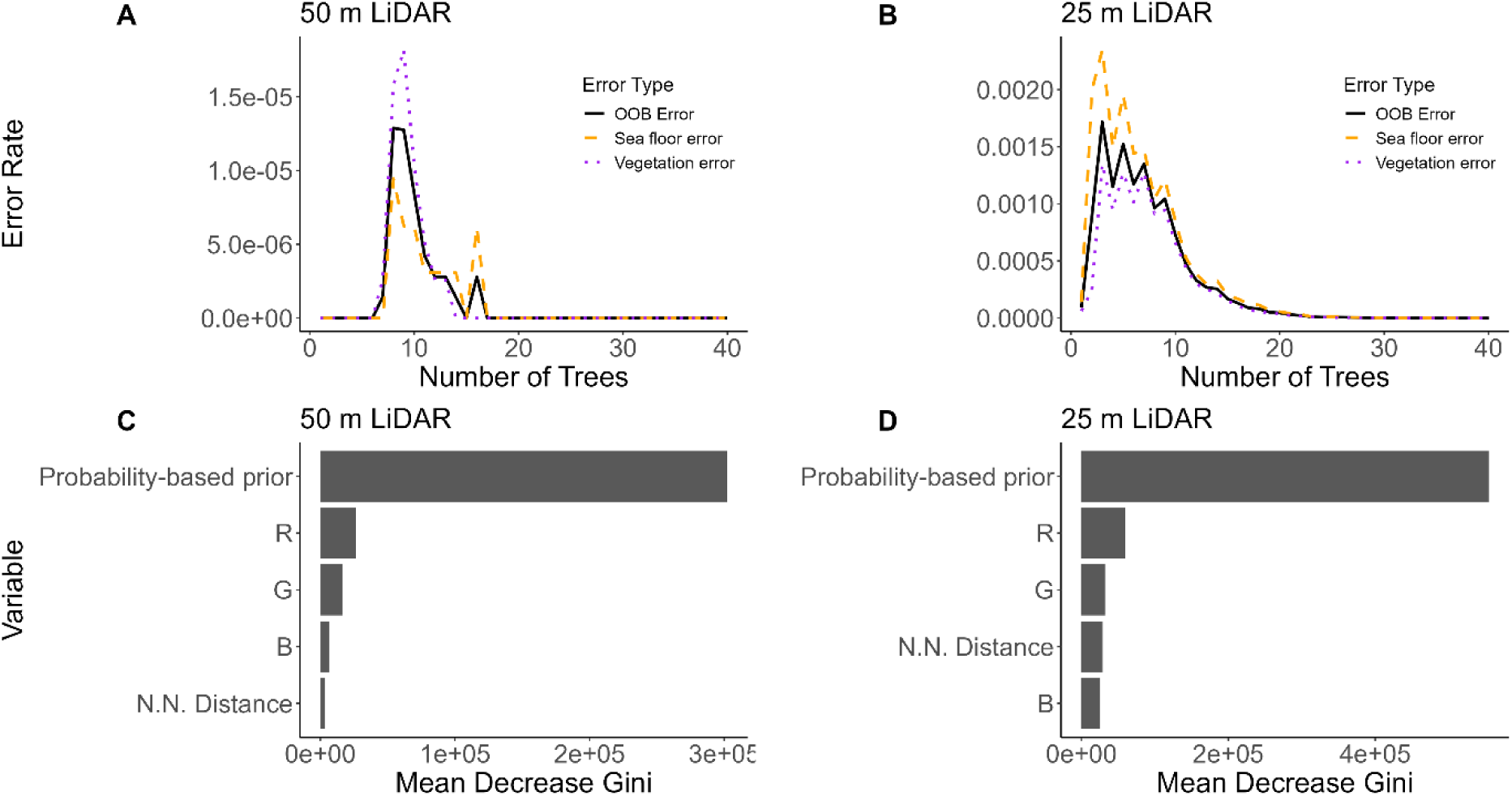
Upper panels shows the Out of Bag (OOB) error rate of the Random Forest model across trees, including the overall rate (black), as well as the rate for the sea floor classification (orange dashed line) and vegetation classification (purple dotted line) for the 50 m (A) and 25 m (B) LiDAR datasets. Lower panels show the relative importance of each variable, measured as Mean Decrease Gini values, within the random forest model, including the vegetation intensity probability-based prior, R, G and B radiometric values, and the nearest neighbour (N.N.) distance of points within the point cloud data for the 50 m (C) and 25 m (D) LiDAR datasets.

**Supplementary Figure 4:**
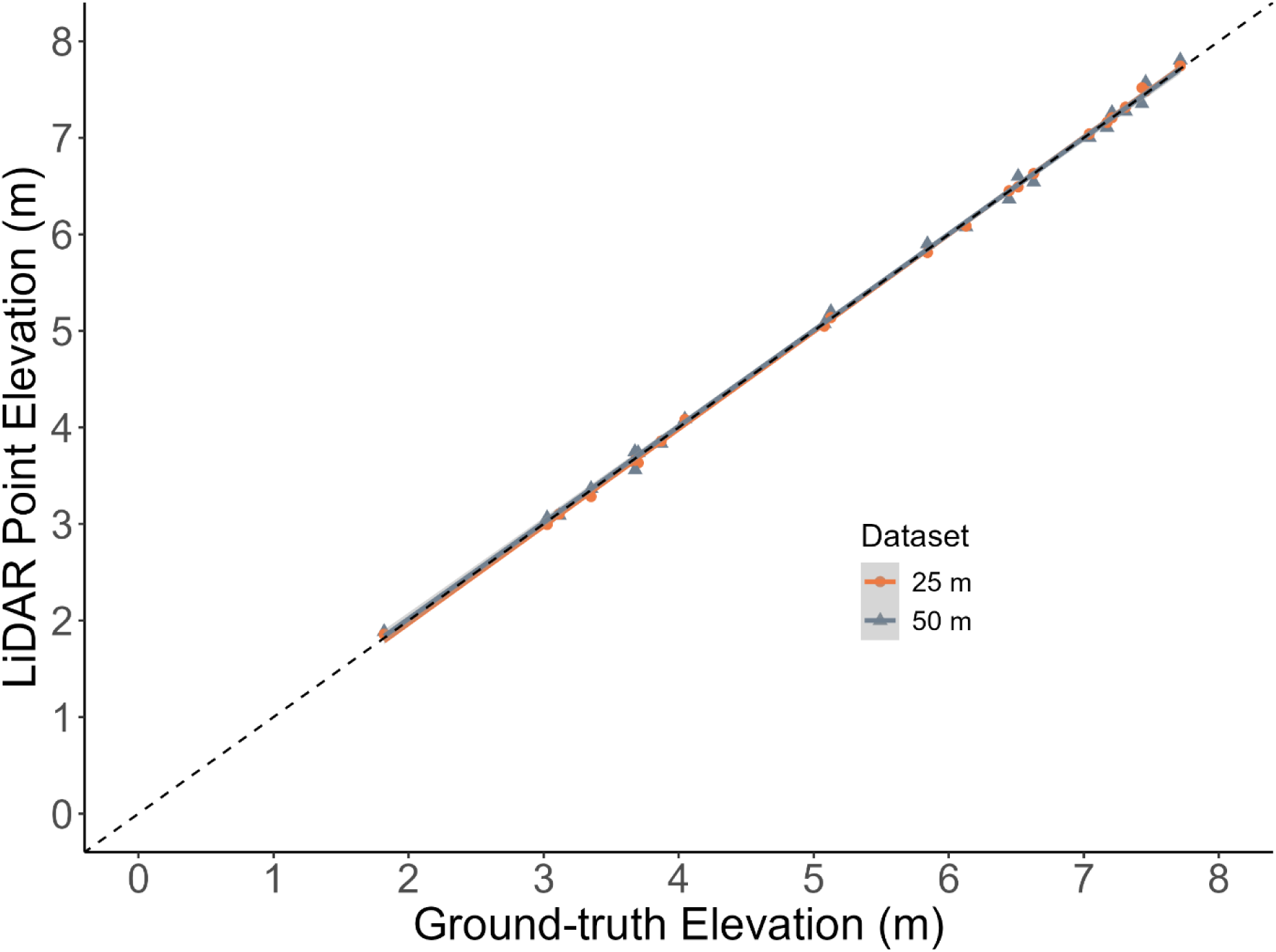
Comparison of above-ground elevation heights between the 25 m LiDAR data (orange) and the 50 m LiDAR data (grey) and handheld positioning Global Navigation Satellite System (GNSS) elevation (m). The dotted line represents a 1:1 relationship.

**Supplementary Figure 5:**
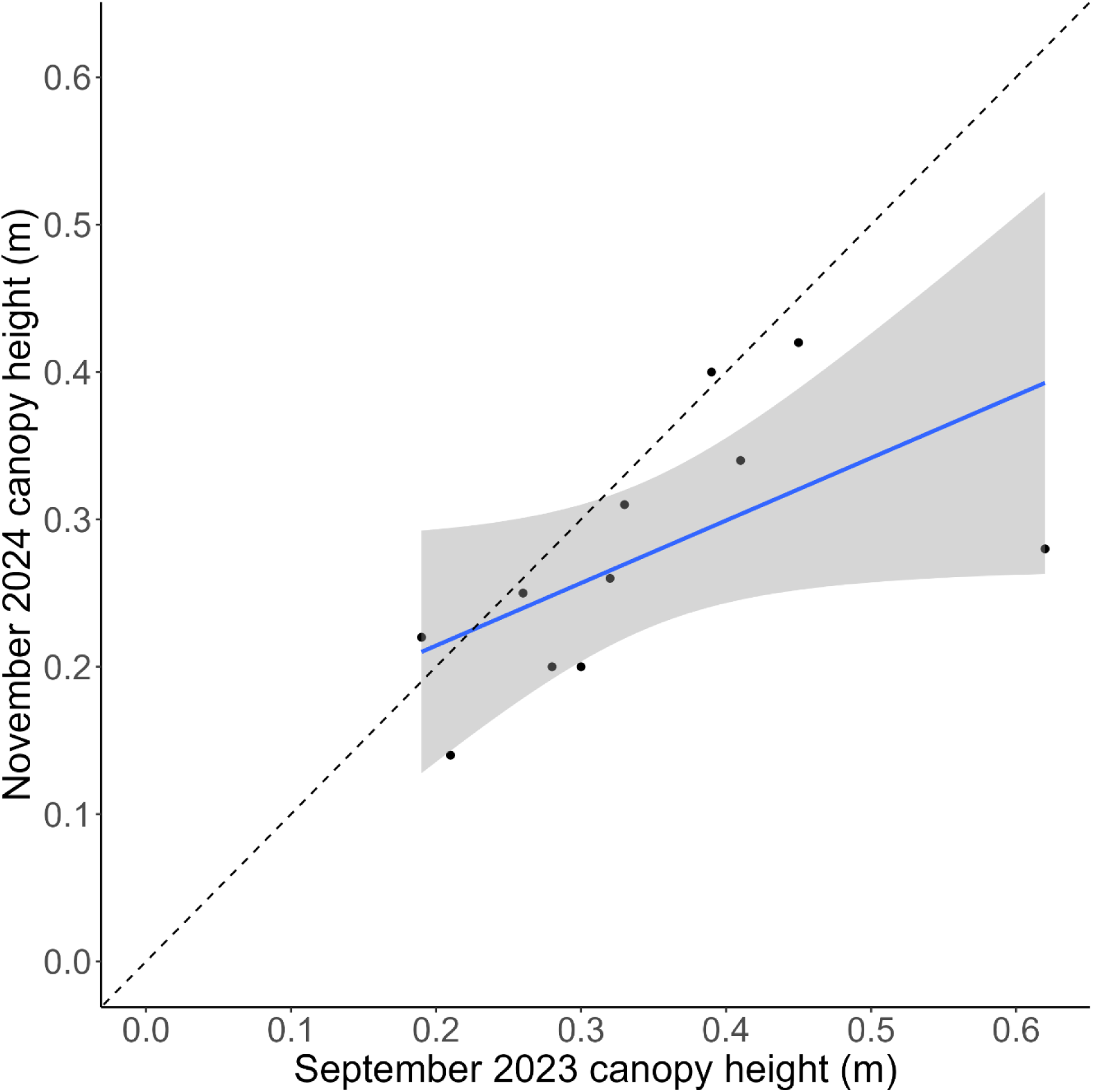
A comparison of eelgrass canopy height (m) measured at the same locations (see Supplementary Figure 1) from two separate field campaigns in September 2023 and November 2024. The dotted line represents a 1:1 relationship.

**Supplementary Figure 6:**
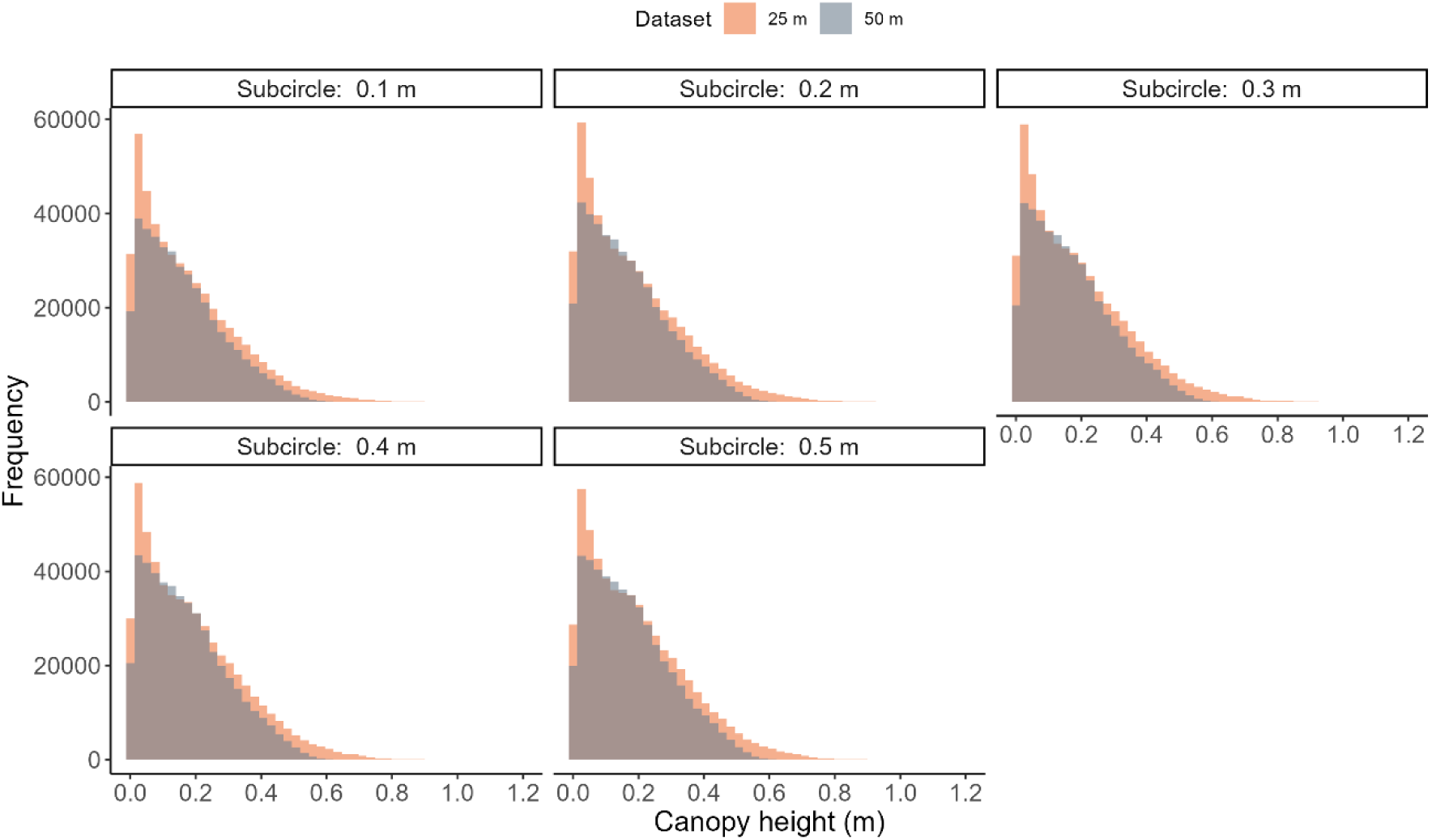
Histograms of canopy height (m) rasters for both the 25 m (orange) and 50 m (grey) LiDAR datasets, across DSM smoothing radii (‘subcircle’ value) from 0.1 to 0.5 m.

**Supplementary Figure 7:**
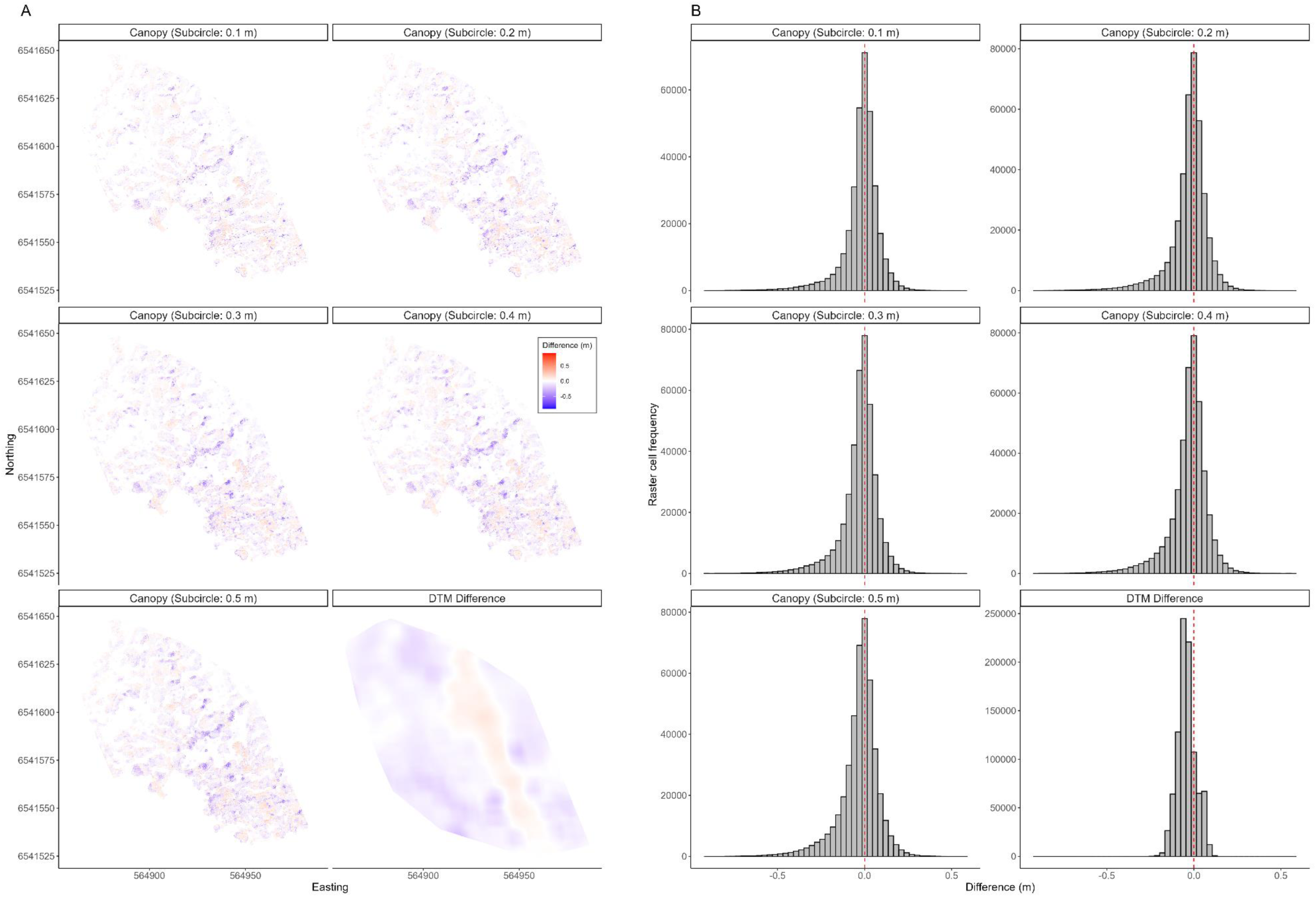
A) Comparison of canopy height rasters (smoothing radii (‘subcircle’ value) from 0.1 – 0.5 m) and sea floor DTM rasters between the 50 m and 25 m LiDAR datasets. The difference (m) between rasters indicates the difference of the 50 m dataset relative to the 25 m dataset, i.e., positive (red) values display higher values for the 50 m dataset and blue values display negative values. B) Histograms of the raster cells displayed in A, displaying the frequency of cells based on their difference (m) values.

**Supplementary Figure 8:**
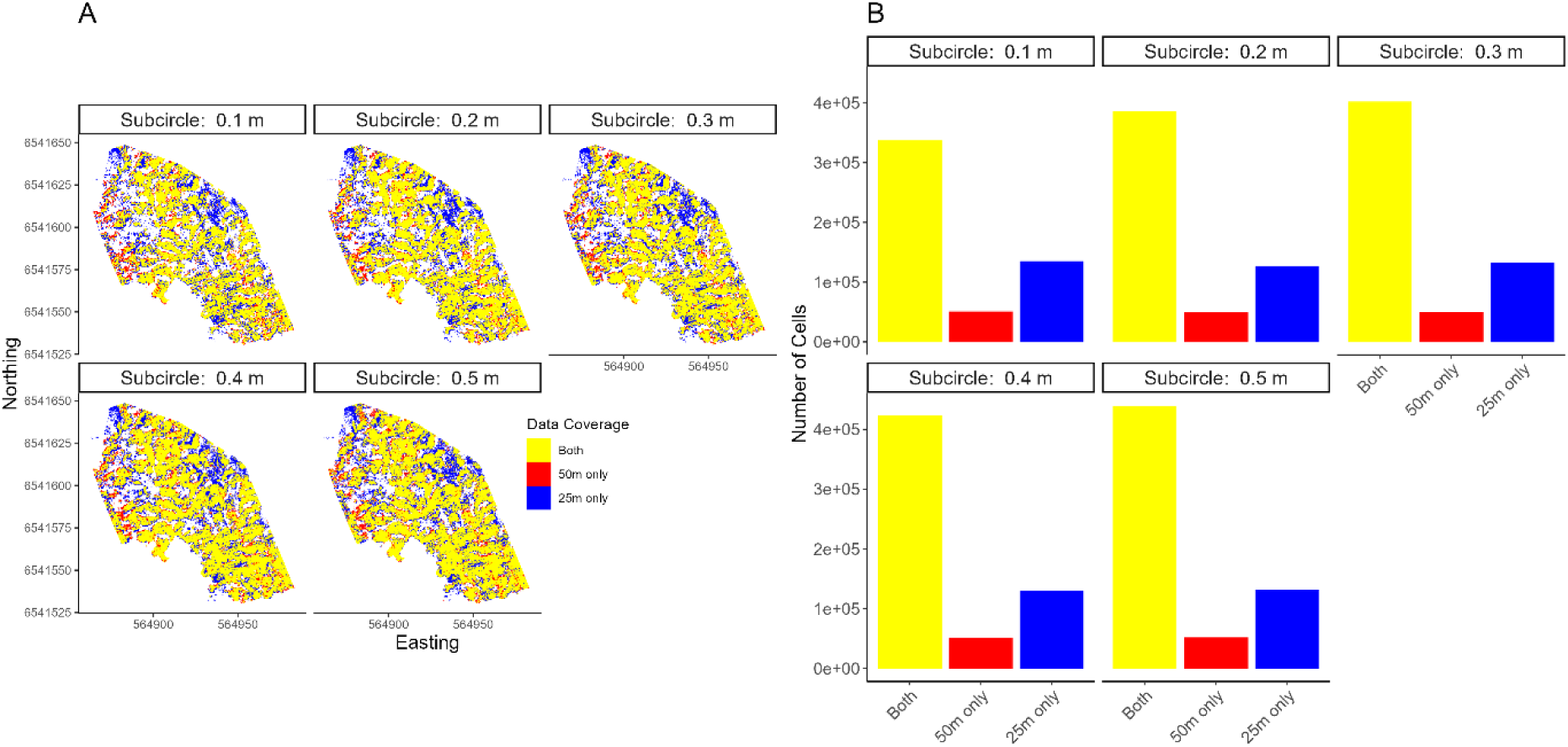
A) Areal coverage of the canopy height rasters between the 50 m (red) and 25 m (blue) datasets, with cells containing values from both datasets displayed in yellow, across the DSM smoothing radii from 0.1 – 0.5 m (‘subcircle’ value). B) A barplot of the raster cells displayed in A, across all smoothing radii, for overlapping cells (both, yellow), the 50 m (red) and the 25 m (blue) dataset.

**Supplementary Table 1:** Eelgrass (*Zostera marina*) samples used to calculate the volumetric biomass, *B_vol_* (g wet weight (ww) m^-3^, see Eq. 3) at the study site in Ølbergholmen, Norway. Samples were collected in September 2023 (see Borger (2024) for details). Measurements taken include eelgrass mean and maximum canopy height (m) within the sample frame area (m^2^). From the volumetric biomass, we converted to dry weight (dw) biomass density (g dw m^-3^) using mean conversion values derived from Supplementary Table 2, and subsequently, to biomass in g Carbon m^-2^ using standard C/dw conversion values (Duarte 1990; Howard and others 2014).

| Height,<br>mean (m) | Height,<br>max (m) | ww (g) | Frame<br>size (cm) | Area ($m^2$) | Biomass<br>g ww $m^{-2}$ | Biomass<br>density<br>(g ww $m^{-3}$) | Biomass<br>(g dw $m^{-2}$) | Biomass<br>density<br>(g dw $m^{-3}$) | Biomass<br>(g C $m^{-2}$) |
| --- | --- | --- | --- | --- | --- | --- | --- | --- | --- |
| 0.30 | 0.50 | 28.71 | 20x20 | 0.04 | 717.74 | 2392.48 | 170.10 | 567.02 | 54.43 |
| 0.20 | 0.30 | 12.74 | 20x20 | 0.04 | 318.53 | 1592.66 | 75.49 | 377.46 | 24.16 |
| 0.18 | 0.40 | 4.48 | 40x40 | 0.04 | 111.95 | 621.96 | 26.53 | 147.40 | 8.49 |
| 0.42 | 0.56 | 6.59 | 40x40 | 0.16 | 41.17 | 98.03 | 9.76 | 23.23 | 3.12 |
| 0.40 | 0.58 | 48.67 | 20x20 | 0.04 | 1216.80 | 3041.99 | 288.38 | 720.95 | 92.28 |
| 0.30 | 0.47 | 10.37 | 20x20 | 0.04 | 259.31 | 864.38 | 61.46 | 204.86 | 19.67 |
| 0.42 | 0.61 | 40.08 | 20x20 | 0.04 | 1001.99 | 2385.68 | 237.47 | 565.41 | 75.99 |
| 0.40 | 0.66 | 54.12 | 20x20 | 0.04 | 1352.95 | 3382.38 | 320.65 | 801.62 | 102.61 |
| 0.32 | 0.57 | 10.69 | 20x20 | 0.04 | 267.26 | 835.20 | 63.34 | 197.94 | 20.27 |
| 0.14 | 0.22 | 1.73 | 20x20 | 0.04 | 43.33 | 309.52 | 10.27 | 73.36 | 3.29 |
| 0.23 | 0.35 | 6.78 | 20x20 | 0.04 | 169.49 | 736.89 | 40.17 | 174.64 | 12.85 |
| 0.26 | 0.41 | 21.28 | 20x20 | 0.04 | 532.09 | 2046.51 | 126.11 | 485.02 | 40.35 |
| 0.16 | 0.28 | 1.00 | 20x20 | 0.04 | 24.88 | 155.47 | 5.90 | 36.85 | 1.89 |
| 0.29 | 0.46 | 37.64 | 20x20 | 0.04 | 941.10 | 3245.18 | 223.04 | 769.11 | 71.37 |
| 0.24 | 0.38 | 16.63 | 20x20 | 0.04 | 415.79 | 1732.47 | 98.54 | 410.60 | 31.53 |
| 0.22 | 0.38 | 4.64 | 20x20 | 0.04 | 116.09 | 527.69 | 27.51 | 125.06 | 8.80 |
| 0.35 | 0.56 | 2.30 | 20x20 | 0.04 | 57.54 | 164.40 | 13.64 | 38.96 | 4.36 |
| 0.30 | 0.57 | 10.64 | 20x20 | 0.04 | 265.96 | 886.53 | 63.03 | 210.11 | 20.17 |
| 0.27 | 0.41 | 4.96 | 20x20 | 0.04 | 123.93 | 458.98 | 29.37 | 108.78 | 9.40 |
| 0.28 | 0.44 | 3.40 | 20x20 | 0.04 | 85.07 | 303.83 | 20.16 | 72.01 | 6.45 |
| 0.21 | 0.45 | 16.03 | 20x20 | 0.04 | 400.77 | 1908.40 | 94.98 | 452.29 | 30.39 |
| 0.19 | 0.35 | 13.94 | 20x20 | 0.04 | 348.44 | 1833.89 | 82.58 | 434.63 | 26.43 |
| 0.31 | 0.49 | 12.20 | 20x20 | 0.04 | 304.95 | 983.72 | 72.27 | 233.14 | 23.13 |
| 0.23 | 0.50 | 17.27 | 20x20 | 0.04 | 431.87 | 1877.67 | 102.35 | 445.01 | 32.75 |
| 0.26 | 0.53 | 9.04 | 20x20 | 0.04 | 226.04 | 869.39 | 53.57 | 206.05 | 17.14 |
| 0.15 | 0.32 | 8.18 | 20x20 | 0.04 | 204.38 | 1362.55 | 48.44 | 322.92 | 15.50 |
| 0.26 | 0.48 | 23.74 | 20x20 | 0.04 | 593.54 | 2282.85 | 140.67 | 541.03 | 45.01 |
| 0.15 | 0.19 | 1.93 | 20x20 | 0.04 | 48.27 | 321.80 | 11.44 | 76.27 | 3.66 |
| 0.15 | 0.28 | 20.68 | 20x20 | 0.04 | 516.99 | 3446.57 | 122.53 | 816.84 | 39.21 |
| 0.13 | 0.33 | 11.35 | 20x20 | 0.04 | 283.69 | 2182.19 | 67.23 | 517.18 | 21.51 |
| 0.19 | 0.43 | 7.37 | 20x20 | 0.04 | 184.37 | 970.38 | 43.70 | 229.98 | 13.98 |
| 0.19 | 0.26 | 5.64 | 20x20 | 0.04 | 140.92 | 741.68 | 33.40 | 175.78 | 10.69 |
| 0.18 | 0.39 | 15.32 | 20x20 | 0.04 | 382.99 | 2127.69 | 90.77 | 504.26 | 29.05 |
| 0.45 | 0.71 | 39.90 | 20x20 | 0.04 | 997.42 | 2216.48 | 236.39 | 525.31 | 75.64 |
| 0.13 | 0.27 | 1.50 | 20x20 | 0.04 | 37.57 | 289.00 | 8.90 | 68.49 | 2.85 |
| 0.21 | 0.32 | 5.22 | 20x20 | 0.04 | 130.50 | 621.44 | 30.93 | 147.28 | 9.90 |
| 0.09 | 0.21 | 1.15 | 20x20 | 0.04 | 28.78 | 319.72 | 6.82 | 75.77 | 2.18 |
| 0.23 | 0.30 | 0.78 | 20x20 | 0.04 | 19.44 | 84.53 | 4.61 | 20.03 | 1.47 |
| 0.36 | 0.62 | 35.69 | 20x20 | 0.04 | 892.24 | 2478.44 | 211.46 | 587.39 | 67.67 |
| 0.22 | 0.41 | 22.97 | 20x20 | 0.04 | 574.31 | 2610.51 | 136.11 | 618.69 | 43.56 |
| 0.20 | 0.30 | 4.73 | 20x20 | 0.04 | 118.15 | 590.75 | 28.00 | 140.01 | 8.96 |

**Supplementary Table 2:** Table calculating mean wet weight to dry weight ratio of eelgrass (*Zostera marina*), from subsamples (*n* = 85) taken from samples gathered at the study area of Ølbergholmen, Norway in September 2023.

| Subsample | Wet weight (g) | Dry weight (g) | Dry:wet weight ratio |
| --- | --- | --- | --- |
| 1 | 1.8851 | 0.4014 | 0.212933001 |
| 2 | 1.8615 | 0.4078 | 0.219070642 |
| 3 | 1.8365 | 0.3921 | 0.213503948 |
| 4 | 2.1094 | 0.4153 | 0.19688063 |
| 5 | 2.8975 | 0.5509 | 0.190129422 |
| 6 | 3.4306 | 0.6543 | 0.190724655 |
| 7 | 2.5313 | 0.522 | 0.206218149 |
| 8 | 1.3271 | 0.2766 | 0.208424384 |
| 9 | 2.2412 | 0.458 | 0.20435481 |
| 10 | 1.5016 | 0.3252 | 0.216568993 |
| 11 | 1.9788 | 0.4905 | 0.247877502 |
| 12 | 1.2281 | 0.2822 | 0.229785848 |
| 13 | 0.6711 | 0.2194 | 0.326925942 |
| 14 | 1.81207 | 0.3483 | 0.192211118 |
| 15 | 2.6198 | 0.4839 | 0.184708756 |
| 16 | 2.8489 | 0.486 | 0.170592158 |
| 17 | 1.4015 | 0.2638 | 0.1882269 |
| 18 | 1.0242 | 0.2241 | 0.218804921 |
| 19 | 1.3848 | 0.2559 | 0.184792028 |
| 20 | 0.7355 | 0.1338 | 0.181917063 |
| 21 | 1.6018 | 0.3204 | 0.200024972 |
| 22 | 1.327 | 0.2172 | 0.163677468 |
| 23 | 1.3902 | 0.2796 | 0.201122141 |
| 24 | 1.7986 | 0.3627 | 0.201656844 |
| 25 | 2.0476 | 0.5259 | 0.256837273 |
| 26 | 2.3109 | 0.4426 | 0.191527111 |
| 27 | 1.4363 | 0.3065 | 0.21339553 |
| 28 | 1.7463 | 0.347 | 0.198705835 |
| 29 | 2.6406 | 0.523 | 0.198061047 |
| 30 | 2.2776 | 0.4704 | 0.206533193 |
| 31 | 1.6274 | 0.3526 | 0.216664618 |
| 32 | 0.8941 | 0.1906 | 0.21317526 |
| 33 | 2.5145 | 0.4582 | 0.182223106 |
| 34 | 1.6931 | 0.3168 | 0.187112397 |
| 35 | 1.1887 | 0.219 | 0.184234878 |
| 36 | 1.8262 | 0.2838 | 0.155404665 |
| 37 | 1.342 | 0.248 | 0.184798808 |
| 38 | 2.2096 | 0.4128 | 0.186821144 |
| 39 | 1.2231 | 0.3216 | 0.262938435 |
| 40 | 2.1034 | 0.4058 | 0.192925739 |
| 41 | 1.7557 | 0.3491 | 0.19883807 |
| 42 | 1.7235 | 0.3043 | 0.176559327 |
| 43 | 2.8747 | 0.5497 | 0.191219953 |
| 44 | 3.0337 | 0.5307 | 0.174934898 |
| 45 | 1.647 | 0.3839 | 0.233090468 |
| 46 | 0.5349 | 0.1655 | 0.309403627 |
| 47 | 0.7314 | 0.1275 | 0.174323216 |
| 48 | 1.01 | 0.3025 | 0.29950495 |
| 49 | 1.162 | 0.2578 | 0.221858864 |
| 50 | 0.8309 | 0.1796 | 0.216151161 |
| 51 | 1.2758 | 0.319 | 0.250039191 |
| 52 | 0.5961 | 0.1591 | 0.266901527 |
| 53 | 0.2702 | 0.0779 | 0.288304959 |
| 54 | 0.4747 | 0.1427 | 0.300610912 |
| 55 | 0.9063 | 0.2271 | 0.250579278 |
| 56 | 0.4232 | 0.0627 | 0.1481569 |
| 57 | 0.442 | 0.1177 | 0.266289593 |
| 58 | 0.6792 | 0.2072 | 0.305064782 |
| 59 | 1.921 | 0.3721 | 0.193701197 |
| 60 | 0.8888 | 0.1524 | 0.171467147 |
| 61 | 0.8221 | 0.2094 | 0.254713538 |
| 62 | 1.2833 | 0.4514 | 0.351749396 |
| 63 | 2.4596 | 0.6047 | 0.245852984 |
| 64 | 0.5653 | 0.1211 | 0.214222537 |
| 65 | 1.6672 | 0.4483 | 0.268893954 |
| 66 | 0.5936 | 0.1843 | 0.310478437 |
| 67 | 1.3044 | 0.3083 | 0.236353879 |
| 68 | 0.7976 | 0.3446 | 0.432046138 |
| 69 | 0.8132 | 0.2995 | 0.368298082 |
| 70 | 1.0338 | 0.2767 | 0.267653318 |
| 71 | 2.5686 | 0.5762 | 0.224324535 |
| 72 | 0.6127 | 0.2582 | 0.421413416 |
| 73 | 1.1841 | 0.2721 | 0.229794781 |
| 74 | 0.7721 | 0.2092 | 0.270949359 |
| 75 | 0.9489 | 0.2637 | 0.277900727 |
| 76 | 0.7381 | 0.2121 | 0.287359436 |
| 77 | 0.857 | 0.2497 | 0.291365228 |
| 78 | 1.0042 | 0.31 | 0.308703446 |
| 79 | 0.4764 | 0.1103 | 0.231528128 |
| 80 | 0.7448 | 0.1889 | 0.253625134 |
| 81 | 0.3233 | 0.0939 | 0.290442314 |
| 82 | 0.3353 | 0.0908 | 0.270802267 |
| 83 | 0.7765 | 0.2279 | 0.293496458 |
| 84 | 0.6892 | 0.1897 | 0.275246663 |
| 85 | 0.8573 | 0.2162 | 0.252187099 |

